# Temporary prey storage along swarm columns of army ants: an adaptive strategy for successful raiding?

**DOI:** 10.1101/2021.05.24.445418

**Authors:** Hilário Póvoas de Lima, Serafino Teseo, Raquel Leite Castro de Lima, Ronara Souza Ferreira-Châline, Nicolas Châline

## Abstract

While pillaging the brood of other ant colonies, *Eciton* army ants accumulate prey in piles, or caches, along their foraging trails. Widely documented, these structures have historically been considered as byproducts of heavy traffic or aborted relocations of the ants’ temporary nest, or bivouac. However, we recently observed that caches of the hook-jawed army ant, *Eciton hamatum*, appeared independently from heavy traffic or bivouac relocations. In addition, the flow of prey through caches varied based on the quantity of prey items workers transported. As this suggested a potential adaptive function, we developed agent-based simulations to compare raids of caching and non-caching virtual army ants. We found that caches increased the amount of prey that relatively low numbers of raiders were able to retrieve. However, this advantage became less conspicuous - and generally disappeared - as the number of raiders increased. Based on these results, we hypothesize that caches maximize the amount of prey that limited amounts of raiders can retrieve, especially as prey colonies coordinately evacuate their brood. In principle, caches also allow workers to safely collect multiple prey items and efficiently transport them to the bivouac. Further field observations are needed to test this and other hypotheses emerging from our study.

## Background

In army ants, foraging occurs through the formation of huge columns of workers roaming forest floors and raiding other social insect colonies [1,2]. During raids, colonies of the hook-jawed army ant, *Eciton hamatum*, accumulate brood prey in caches along their columns (Figure 1A). Pioneer army ant scientists attributed this behavior to traffic management inefficiencies. Schneirla, for example, noticed that numerous workers swarming from the bivouac towards the foraging fronts prevented prey-carrying foragers from returning, ‘virtually forcing’ them ‘to deposit their burdens in piles that form near the places of greatest confusion’ [3]. Rettenmeyer later suggested that caches emerge as prey-carrying workers gather in ‘areas of greater booty odor’, eventually leading to the formation of new bivouacs if caches become especially large [4]. His observations implied that caches are by-products of bivouac regular relocation.

**Figure 1.**
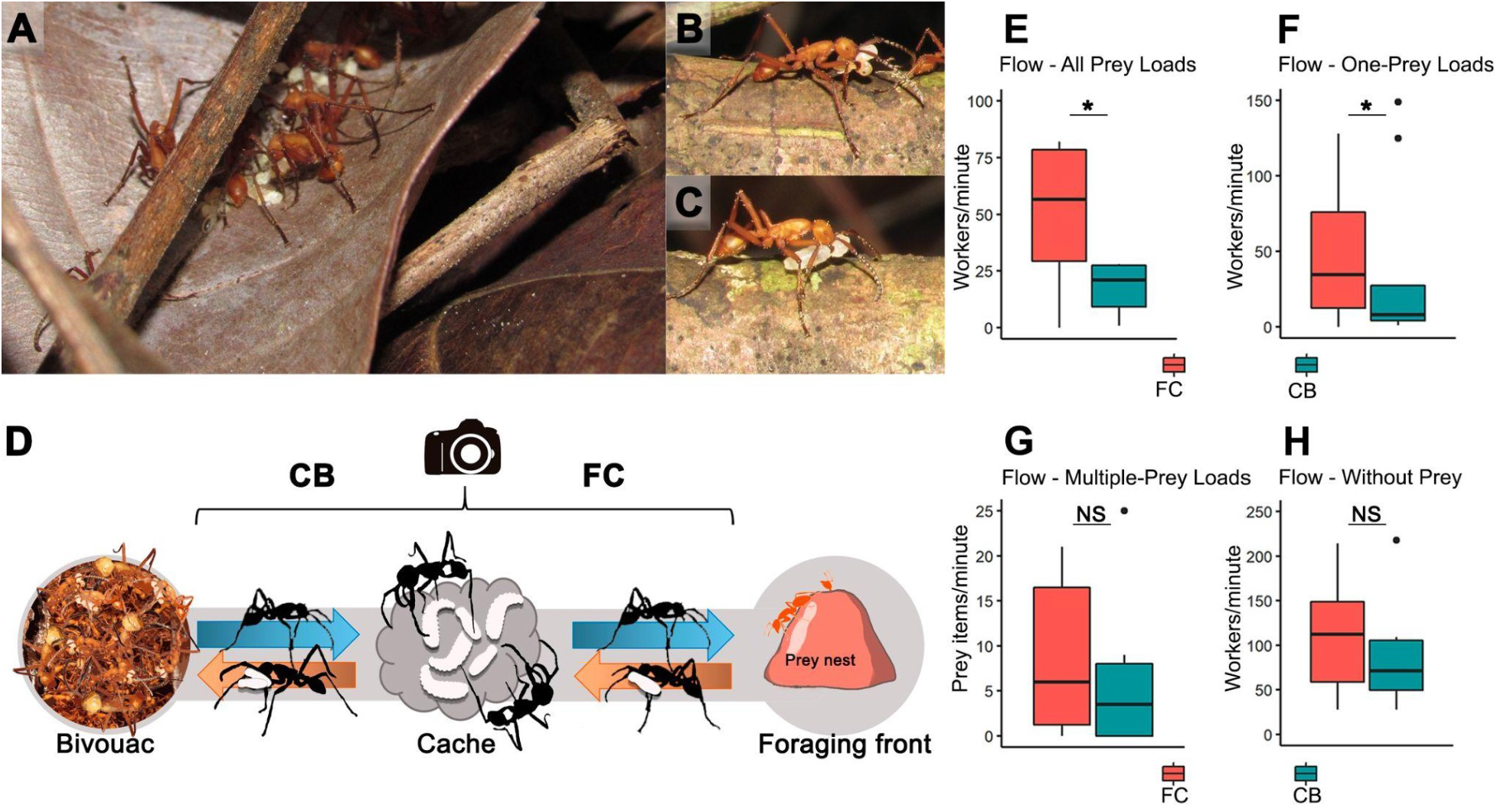
**A.** *Eciton hamatum* cache. **B.** Worker carrying two prey items. **C.** Worker carrying one prey item. **D.** Scheme of field observations. **E**. Flow of all prey-carrying workers. **F**. Workers carrying one prey item. **G**. Workers carrying multiple prey items. **H**. Workers without prey. In whisker plots, central lines, boxes, whiskers and dots respectively show median, quartiles, max/min and outliers. ***: p < 0.001, **: p < 0.01, *: p < 0.05; NS: no significance.

Observing the foraging activity of *E. hamatum*, we noticed that caches appeared regularly even at low traffic intensities and at times of the day in which colonies do not usually relocate, raising doubts about the hypothesis that caches exclusively emerge as byproducts. This idea was corroborated by other experimental work on *Atta* leaf-cutting ants, which also transport huge food quantities along long trails, showing that leaf fragment caches emerge at nest entrances when food inflow exceeds processing rates, and reduce the costs of vertical transport [5,6]. As *Atta* workers maximize food collection via unloading at caches and rapidly resuming foraging, we hypothesized that *E. hamatum* caches may similarly serve to maximize prey retrieval. Therefore, after measuring ant traffic and prey transport through caches in natural conditions, we explored this hypothesis using agent-based simulations, aiming to determine whether and how caches provide a selective advantage.

## Results and discussion

Prior to field observations, we defined caches as structures including stacked prey brood, stationary *E. hamatum* workers (Figure 1A, B, C, video S1) and approaching/leaving individuals. Then, following foraging columns, we found ten caches, six of which included 116±130.56 prey items (total=697; min=18; max=296; all ants, mainly *Pheidole* and *Linepithema*; Figure S1, Tables S1, S2; Supplementary Material text). We found no bivouacs in the surrounding 10m radius. As we conducted observations between 8:00 am and 4:30 pm, we concluded that the observed caches did not originate from traffic bottlenecks or aborted bivouacs, which instead emerge immediately before sunset [3].

Analyzing videos of ant traffic through caches (Figure 1D), we counted 189.8±117.4 workers going from the bivouac to the foraging fronts and 226.1±116.3 workers in the opposite direction, transporting 75±71.9 prey items. We found no significant differences between the numbers of workers in the video frame portion including the stretch between the foraging front and the cache (FC), and that between the cache and the bivouac (CB), for individuals traveling from the foraging front to the bivouac (t=16, p=0.24), in the opposite direction (t=27, p=0.95) and in both directions pooled (U= 33.50, p=0.21). At caches, we recorded more prey loads passing through FC than CB (t= 6.0; p = 0.05, Figure 1E) and more workers carrying single-item prey loads in FC than CB (57.2±63.2 vs. 35.6±54.5; t=8.0; n=10, p=0.04, Figure 1F). The numbers of workers carrying multiple-item prey loads did not differ significantly (FC: 8.9±8.8; BC: 7.6±10.9; t=13.5; p=0.52, Figure 1G). This indicated that, at caches, ants accumulated single-prey loads, whereas multiple-prey loads, although arriving at caches in significantly lower numbers (8.9±8.8 compared to 57.2±63.2 single-prey loads, U = 75.5, p < 0.05), flowed relatively regularly. We found no differences between the numbers of unloaded workers in FC and BC (151.4±161.9 vs. 116.5±120.8; t=11; p=0.09, Figure 1H). If caches only emerged as by-products, we would not expect prey accumulation/transport to depend on load size. We therefore suspected that, similar to the caches of leaf cutting ants [5,6], *E. hamatum*’s caches may serve to optimize colony-level foraging investments. We hypothesized that caches may emerge as raiders returning from foraging fronts drop single-prey loads in safe locations, and rapidly return to foraging fronts. From a colony-level perspective, short sequential trips between caches and foraging fronts would maximize prey yields at limited numbers of raiders, especially because prey colonies coordinately evacuate their brood [7,8]. In addition, in the chaos of raids, rapidly retrieving single prey items may be safer and more convenient than sequentially collecting multiple prey items. On the other hand, workers at caches could invest time in loading multiple items in a significantly safer microenvironment, minimizing the distance walked per retrieved food mass on their way to the bivouac.

To explore such hypotheses, and investigate whether and in which conditions caches would increase prey retrieval, we developed Netlogo agent-based simulations (Video S2, Table S3, Supplementary Material text) [9]. Our virtual ants formed columns via releasing/following a trail pheromone, encountering prey item piles simulating the brood of prey colonies. As ants began raiding, uncollected prey items started disappearing, simulating brood evacuation. Raiders collected prey items and cached them with a probability increasing with nestmate density, simulating the scenario we observed in nature. Other workers recovered up to two cached prey items and transported them to a densely populated “safe area” simulating a trail bifurcation or the bivouac itself. When loaded raiders or cache recoverers reached the safe area, prey items disappeared and were counted as retrieved. We compared caching colonies to non-caching colonies in which raiders transported prey items directly to the safe area.

Overall, the quantity of retrieved prey increased with the number of raiders, but decreased as brood evacuation speeds and probability to cache/collect cached food increased (Table 1, Model 1). Interestingly, caches allowed ants to retrieve significantly more prey at low numbers of raiders (50, 100), but this effect became generally less conspicuous - and disappeared - as raider numbers increased (300, 500, 700; Figure 2, Table 1, Models 2-5). Brood evacuation speed reduced the amount of prey ants collected at foraging fronts, in turn reducing the amount of time ants spent retrieving it. Excluding the 700-raider condition, the time ants spent retrieving prey decreased at increasing raider numbers, whereas the probability of caching/collecting cached prey did not produce any effect (Table 1, Model 6). Hardly any combination of settings revealed significant effects of caches on prey retrieval time (Table S4, Figure S2), indicating that, everything else being equal, caching ants retrieved food at the same speed of non-caching ones. At very high raider numbers, however, the high individual density occasionally resulted in persisting ‘death circles’, decelerating prey retrieval. Allowing cache recoverers to keep caching their loads after collection rarely produced significant effects on the amount of retrieved prey (Table S5, Figure S3), but increased prey retrieval time at very low and very high numbers of raiders (Table S6, Figure S4). This likely occurred because raiders in 50-100-individual groups encountered cached food items relatively rarely, whereas 500-700-raider groups kept encountering high densities of individuals, caching food very often. This suggested that excessive caching may slow down prey retrieval, and accordingly, our field observations revealed a relatively stable flow of multiple-item prey loads through caches (Figure 1G). Therefore, we hypothesize that, for cache recoverers transporting multiple prey items, it may be advantageous to unload only at the bivouac and not in other caches.

**Figure 2.**
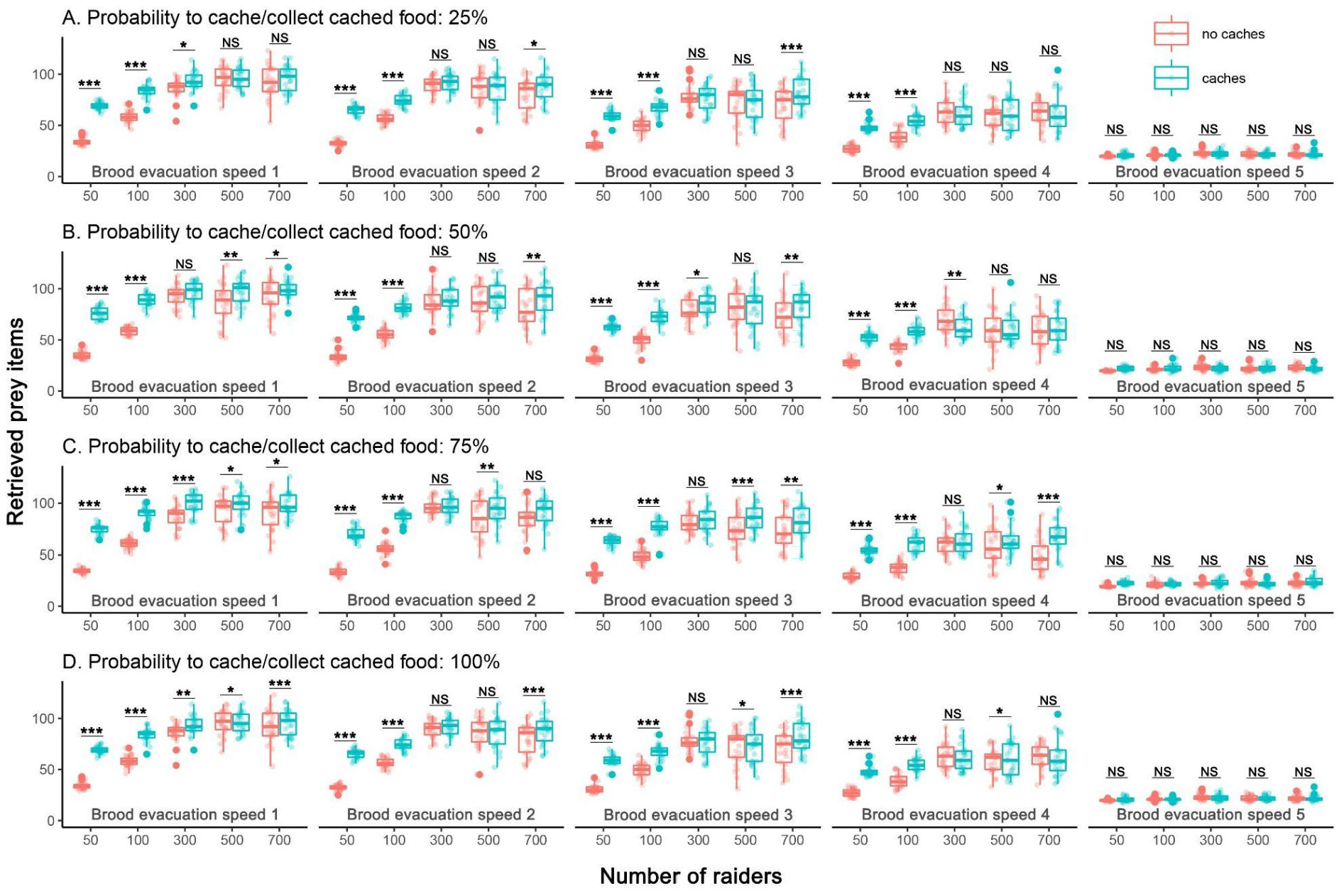
Simulation results. Amount of retrieved prey. In whisker plots (each representing 25 simulations), central lines, boxes, whiskers and dots respectively show median, quartiles, max/min and outliers. ***: p < 0.001, **: p < 0.01, *: p < 0.05; NS: no significance.

**Table 1.**
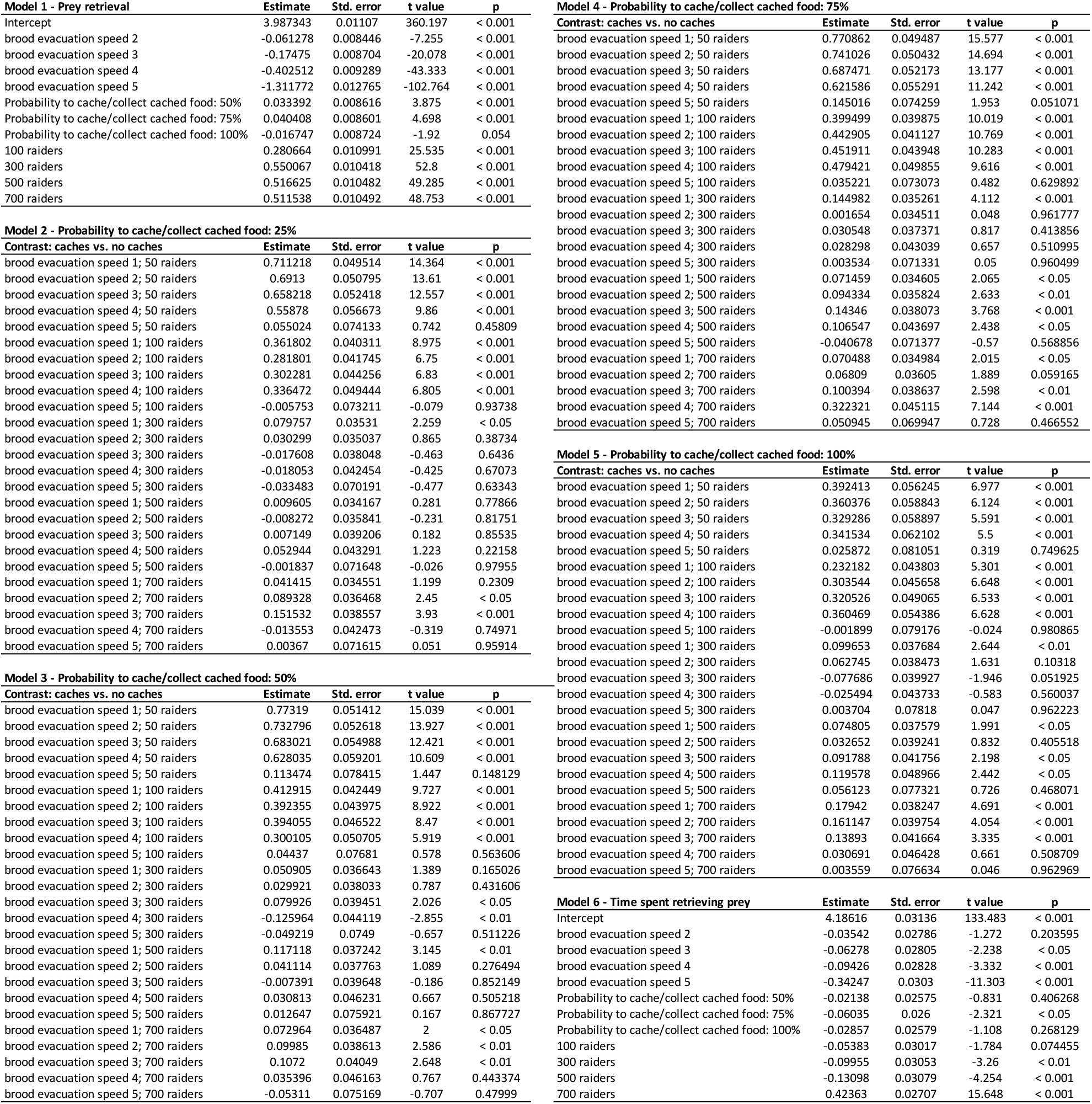
Models’ outputs.

| Model 1 - Prey retrieval | Estimate | Std. error | t value | p |
| --- | --- | --- | --- | --- |
| Intercept | 3.987343 | 0.01107 | 360.197 | < 0.001 |
| brood evacuation speed 2 | -0.061278 | 0.008446 | -7.255 | < 0.001 |
| brood evacuation speed 3 | -0.17475 | 0.008704 | -20.078 | < 0.001 |
| brood evacuation speed 4 | -0.402512 | 0.009289 | -43.333 | < 0.001 |
| brood evacuation speed 5 | -1.311772 | 0.012765 | -102.764 | < 0.001 |
| Probability to cache/collect cached food: 50% | 0.033392 | 0.008616 | 3.875 | < 0.001 |
| Probability to cache/collect cached food: 75% | 0.040408 | 0.008601 | 4.698 | < 0.001 |
| Probability to cache/collect cached food: 100% | -0.016747 | 0.008724 | -1.92 | 0.054 |
| 100 raiders | 0.280664 | 0.010991 | 25.535 | < 0.001 |
| 300 raiders | 0.550067 | 0.010418 | 52.8 | < 0.001 |
| 500 raiders | 0.516625 | 0.010482 | 49.285 | < 0.001 |
| 700 raiders | 0.511538 | 0.010492 | 48.753 | < 0.001 |

| Model 2 - Probability to cache/collect cached food: 25% | Estimate | Std. error | t value | p |
| --- | --- | --- | --- | --- |
| Contrast: caches vs. no caches |  |  |  |  |
| brood evacuation speed 1; 50 raiders | 0.711218 | 0.049514 | 14.364 | < 0.001 |
| brood evacuation speed 2; 50 raiders | 0.6913 | 0.050795 | 13.61 | < 0.001 |
| brood evacuation speed 3; 50 raiders | 0.658218 | 0.052418 | 12.557 | < 0.001 |
| brood evacuation speed 4; 50 raiders | 0.55878 | 0.056673 | 9.86 | < 0.001 |
| brood evacuation speed 5; 50 raiders | 0.055024 | 0.074133 | 0.742 | 0.45809 |
| brood evacuation speed 1; 100 raiders | 0.361802 | 0.040311 | 8.975 | < 0.001 |
| brood evacuation speed 2; 100 raiders | 0.281801 | 0.041745 | 6.75 | < 0.001 |
| brood evacuation speed 3; 100 raiders | 0.302281 | 0.044256 | 6.83 | < 0.001 |
| brood evacuation speed 4; 100 raiders | 0.336472 | 0.049444 | 6.805 | < 0.001 |
| brood evacuation speed 5; 100 raiders | -0.005753 | 0.073211 | -0.079 | 0.93738 |
| brood evacuation speed 1; 300 raiders | 0.079757 | 0.03531 | 2.259 | < 0.05 |
| brood evacuation speed 2; 300 raiders | 0.030299 | 0.035037 | 0.865 | 0.38734 |
| brood evacuation speed 3; 300 raiders | -0.017608 | 0.038048 | -0.463 | 0.6436 |
| brood evacuation speed 4; 300 raiders | -0.018053 | 0.042454 | -0.425 | 0.67073 |
| brood evacuation speed 5; 300 raiders | -0.033483 | 0.070191 | -0.477 | 0.63343 |
| brood evacuation speed 1; 500 raiders | 0.009605 | 0.034167 | 0.281 | 0.77866 |
| brood evacuation speed 2; 500 raiders | -0.008272 | 0.035841 | -0.231 | 0.81751 |
| brood evacuation speed 3; 500 raiders | 0.007149 | 0.039206 | 0.182 | 0.85535 |
| brood evacuation speed 4; 500 raiders | 0.052944 | 0.043291 | 1.223 | 0.22158 |
| brood evacuation speed 5; 500 raiders | -0.001837 | 0.071648 | -0.026 | 0.97955 |
| brood evacuation speed 1; 700 raiders | 0.041415 | 0.034551 | 1.199 | 0.2309 |
| brood evacuation speed 2; 700 raiders | 0.089328 | 0.036468 | 2.45 | < 0.05 |
| brood evacuation speed 3; 700 raiders | 0.151532 | 0.038557 | 3.93 | < 0.001 |
| brood evacuation speed 4; 700 raiders | -0.013553 | 0.042473 | -0.319 | 0.74971 |
| brood evacuation speed 5; 700 raiders | 0.00367 | 0.071615 | 0.051 | 0.95914 |

| Model 3 - Probability to cache/collect cached food: 50% | Estimate | Std. error | t value | p |
| --- | --- | --- | --- | --- |
| Contrast: caches vs. no caches |  |  |  |  |
| brood evacuation speed 1; 50 raiders | 0.77319 | 0.051412 | 15.039 | < 0.001 |
| brood evacuation speed 2; 50 raiders | 0.732796 | 0.052618 | 13.927 | < 0.001 |
| brood evacuation speed 3; 50 raiders | 0.683021 | 0.054988 | 12.421 | < 0.001 |
| brood evacuation speed 4; 50 raiders | 0.628035 | 0.059201 | 10.609 | < 0.001 |
| brood evacuation speed 5; 50 raiders | 0.113474 | 0.078415 | 1.447 | 0.148129 |
| brood evacuation speed 1; 100 raiders | 0.412915 | 0.042449 | 9.727 | < 0.001 |
| brood evacuation speed 2; 100 raiders | 0.392355 | 0.043975 | 8.922 | < 0.001 |
| brood evacuation speed 3; 100 raiders | 0.394055 | 0.046522 | 8.47 | < 0.001 |
| brood evacuation speed 4; 100 raiders | 0.300105 | 0.050705 | 5.919 | < 0.001 |
| brood evacuation speed 5; 100 raiders | 0.04437 | 0.07681 | 0.578 | 0.563606 |
| brood evacuation speed 1; 300 raiders | 0.050905 | 0.036643 | 1.389 | 0.165026 |
| brood evacuation speed 2; 300 raiders | 0.029921 | 0.038033 | 0.787 | 0.431606 |
| brood evacuation speed 3; 300 raiders | 0.079926 | 0.039451 | 2.026 | < 0.05 |
| brood evacuation speed 4; 300 raiders | -0.125964 | 0.044119 | -2.855 | < 0.01 |
| brood evacuation speed 5; 300 raiders | -0.049219 | 0.0749 | -0.657 | 0.511226 |
| brood evacuation speed 1; 500 raiders | 0.117118 | 0.037242 | 3.145 | < 0.01 |
| brood evacuation speed 2; 500 raiders | 0.041114 | 0.037763 | 1.089 | 0.276494 |
| brood evacuation speed 3; 500 raiders | -0.007391 | 0.039648 | -0.186 | 0.852149 |
| brood evacuation speed 4; 500 raiders | 0.030813 | 0.046231 | 0.667 | 0.505218 |
| brood evacuation speed 5; 500 raiders | 0.012647 | 0.075921 | 0.167 | 0.867727 |
| brood evacuation speed 1; 700 raiders | 0.072964 | 0.036487 | 2 | < 0.05 |
| brood evacuation speed 2; 700 raiders | 0.09985 | 0.038613 | 2.586 | < 0.01 |
| brood evacuation speed 3; 700 raiders | 0.1072 | 0.04049 | 2.648 | < 0.01 |
| brood evacuation speed 4; 700 raiders | 0.035396 | 0.046163 | 0.767 | 0.443374 |
| brood evacuation speed 5; 700 raiders | -0.05311 | 0.075169 | -0.707 | 0.47999 |

| Model 4 - Probability to cache/collect cached food: 75% | Estimate | Std. error | t value | p |
| --- | --- | --- | --- | --- |
| Contrast: caches vs. no caches |  |  |  |  |
| brood evacuation speed 1; 50 raiders | 0.770862 | 0.049487 | 15.577 | < 0.001 |
| brood evacuation speed 2; 50 raiders | 0.741026 | 0.050432 | 14.694 | < 0.001 |
| brood evacuation speed 3; 50 raiders | 0.687471 | 0.052173 | 13.177 | < 0.001 |
| brood evacuation speed 4; 50 raiders | 0.621586 | 0.055291 | 11.242 | < 0.001 |
| brood evacuation speed 5; 50 raiders | 0.145016 | 0.074259 | 1.953 | 0.051071 |
| brood evacuation speed 1; 100 raiders | 0.399499 | 0.039875 | 10.019 | < 0.001 |
| brood evacuation speed 2; 100 raiders | 0.442905 | 0.041127 | 10.769 | < 0.001 |
| brood evacuation speed 3; 100 raiders | 0.451911 | 0.043948 | 10.283 | < 0.001 |
| brood evacuation speed 4; 100 raiders | 0.479421 | 0.049855 | 9.616 | < 0.001 |
| brood evacuation speed 5; 100 raiders | 0.035221 | 0.073073 | 0.482 | 0.629892 |
| brood evacuation speed 1; 300 raiders | 0.144982 | 0.035261 | 4.112 | < 0.001 |
| brood evacuation speed 2; 300 raiders | 0.001654 | 0.034511 | 0.048 | 0.961777 |
| brood evacuation speed 3; 300 raiders | 0.030548 | 0.037371 | 0.817 | 0.413856 |
| brood evacuation speed 4; 300 raiders | 0.028298 | 0.043039 | 0.657 | 0.510995 |
| brood evacuation speed 5; 300 raiders | 0.003534 | 0.071331 | 0.05 | 0.960499 |
| brood evacuation speed 1; 500 raiders | 0.071459 | 0.034605 | 2.065 | < 0.05 |
| brood evacuation speed 2; 500 raiders | 0.094334 | 0.035824 | 2.633 | < 0.01 |
| brood evacuation speed 3; 500 raiders | 0.14346 | 0.038073 | 3.768 | < 0.001 |
| brood evacuation speed 4; 500 raiders | 0.106547 | 0.043697 | 2.438 | < 0.05 |
| brood evacuation speed 5; 500 raiders | -0.040678 | 0.071377 | -0.57 | 0.568856 |
| brood evacuation speed 1; 700 raiders | 0.070488 | 0.034984 | 2.015 | < 0.05 |
| brood evacuation speed 2; 700 raiders | 0.06809 | 0.03605 | 1.889 | 0.059165 |
| brood evacuation speed 3; 700 raiders | 0.100394 | 0.038637 | 2.598 | < 0.01 |
| brood evacuation speed 4; 700 raiders | 0.322321 | 0.045115 | 7.144 | < 0.001 |
| brood evacuation speed 5; 700 raiders | 0.050945 | 0.069947 | 0.728 | 0.466552 |

| Model 5 - Probability to cache/collect cached food: 100% | Estimate | Std. error | t value | p |
| --- | --- | --- | --- | --- |
| Contrast: caches vs. no caches |  |  |  |  |
| brood evacuation speed 1; 50 raiders | 0.392413 | 0.056245 | 6.977 | < 0.001 |
| brood evacuation speed 2; 50 raiders | 0.360376 | 0.058843 | 6.124 | < 0.001 |
| brood evacuation speed 3; 50 raiders | 0.329286 | 0.058897 | 5.591 | < 0.001 |
| brood evacuation speed 4; 50 raiders | 0.341534 | 0.062102 | 5.5 | < 0.001 |
| brood evacuation speed 5; 50 raiders | 0.025872 | 0.081051 | 0.319 | 0.749625 |
| brood evacuation speed 1; 100 raiders | 0.232182 | 0.043803 | 5.301 | < 0.001 |
| brood evacuation speed 2; 100 raiders | 0.303544 | 0.045658 | 6.648 | < 0.001 |
| brood evacuation speed 3; 100 raiders | 0.320526 | 0.049065 | 6.533 | < 0.001 |
| brood evacuation speed 4; 100 raiders | 0.360469 | 0.054386 | 6.628 | < 0.001 |
| brood evacuation speed 5; 100 raiders | -0.001899 | 0.079176 | -0.024 | 0.980865 |
| brood evacuation speed 1; 300 raiders | 0.099653 | 0.037684 | 2.644 | < 0.01 |
| brood evacuation speed 2; 300 raiders | 0.062745 | 0.038473 | 1.631 | 0.10318 |
| brood evacuation speed 3; 300 raiders | -0.077686 | 0.039927 | -1.946 | 0.051925 |
| brood evacuation speed 4; 300 raiders | -0.025494 | 0.043733 | -0.583 | 0.560037 |
| brood evacuation speed 5; 300 raiders | 0.003704 | 0.07818 | 0.047 | 0.962223 |
| brood evacuation speed 1; 500 raiders | 0.074805 | 0.037579 | 1.991 | < 0.05 |
| brood evacuation speed 2; 500 raiders | 0.032652 | 0.039241 | 0.832 | 0.405518 |
| brood evacuation speed 3; 500 raiders | 0.091788 | 0.041756 | 2.198 | < 0.05 |
| brood evacuation speed 4; 500 raiders | 0.119578 | 0.048966 | 2.442 | < 0.05 |
| brood evacuation speed 5; 500 raiders | 0.056123 | 0.077321 | 0.726 | 0.468071 |
| brood evacuation speed 1; 700 raiders | 0.17942 | 0.038247 | 4.691 | < 0.001 |
| brood evacuation speed 2; 700 raiders | 0.161147 | 0.039754 | 4.054 | < 0.001 |
| brood evacuation speed 3; 700 raiders | 0.13893 | 0.041664 | 3.335 | < 0.001 |
| brood evacuation speed 4; 700 raiders | 0.030691 | 0.046428 | 0.661 | 0.508709 |
| brood evacuation speed 5; 700 raiders | 0.003559 | 0.076634 | 0.046 | 0.962969 |

| Model 6 - Time spent retrieving prey | Estimate | Std. error | t value | p |
| --- | --- | --- | --- | --- |
| Intercept | 4.18616 | 0.03136 | 133.483 | < 0.001 |
| brood evacuation speed 2 | -0.03542 | 0.02786 | -1.272 | 0.203595 |
| brood evacuation speed 3 | -0.06278 | 0.02805 | -2.238 | < 0.05 |
| brood evacuation speed 4 | -0.09426 | 0.02828 | -3.332 | < 0.001 |
| brood evacuation speed 5 | -0.34247 | 0.0303 | -11.303 | < 0.001 |
| Probability to cache/collect cached food: 50% | -0.02138 | 0.02575 | -0.831 | 0.406268 |
| Probability to cache/collect cached food: 75% | -0.06035 | 0.026 | -2.321 | < 0.05 |
| Probability to cache/collect cached food: 100% | -0.02857 | 0.02579 | -1.108 | 0.268129 |
| 100 raiders | -0.05383 | 0.03017 | -1.784 | 0.074455 |
| 300 raiders | -0.09955 | 0.03053 | -3.26 | < 0.01 |
| 500 raiders | -0.13098 | 0.03079 | -4.254 | < 0.001 |
| 700 raiders | 0.42363 | 0.02707 | 15.648 | < 0.001 |

Overall, our findings are in line with the hypothesis that *E. hamatum* caches do not only emerge as heavy traffic byproducts or as aborted bivouacs, but also increase prey collection efficiency. They may maximize prey retrieval at limited raider numbers - suggesting raider availability as a factor limiting prey collection at foraging fronts - and when attacked prey colonies coordinate to evacuate their brood. Importantly, caches may rule out the need to recruit large numbers of workers, crucially saving the time prey colonies need to evacuate their brood.

Further potential research questions concern caching from proximal and individual perspectives. Experience affects ant behavioral ontogeny and task partitioning [10], but we ignore its impact in large, complex societies. For example, do *Eciton* foragers specialize in raiding at foraging fronts or in commuting between caches and the bivouac? Similarly, shortterm experience at foraging fronts (e.g., nest/prey features, prey colony defenses) or at caches (e.g., number/type of prey items [7,8]) may affect individual foraging decisions. Caches may allow transfer of information about prey colonies [11,12] and traffic intensity, reducing time-consuming, risky, unnecessary travel. We also ignore whether returning raiders stop at caches or proceed depending on prey load size. A potential proximal cause of this would be the stimulus originating from the extension of the mandibles, greater extension meaning heavier and more cumbersome loads. Another possibility is experience or age-dependent polyethism relegating younger workers to traveling between foraging fronts and prey caches, with older individuals specializing in raiding in a classic task partitioning paradigm [13]. Whatever the mechanism, an ultimate cause explanation is that individuals carrying multiple prey items should proceed straight to the bivouac, saving the unloading time and the time for other workers to further load/unload multiple prey items. The relatively stable flow of multiple-item prey loads we observed at caches supports this hypothesis.

Prey individual/colony size and specific defense strategies also potentially affect cache emergence. In our study, small-sized *Pheidole* and *Linepithema* ant brood dominated cached prey composition, possibly because caching single prey items and transporting them to the bivouac in multiple-item loads is more efficient for tiny than large-sized prey. Future long-term sampling across habitats/seasons can reveal whether *E. hamatum* iteratively adjusts its raiding strategies at a local scale in a prey-dependent fashion. Finally, caches may contribute to safely storing prey when returning directly to the bivouac is risky. For example, in case of rain, stocking prey under the leaf litter may increase chances to successfully transport it to the bivouac at a later time.

## Conclusions

Food caches related to traffic partitioning emerge across distant ant taxa (i.e *Atta, Camponotus* and *Eciton* [2,5,6,14–18], usually in societies where thousands of individuals transport large amounts of food through long distances [19–21]. These must carefully balance foraging investments in terms of energy and time [22–24]. In this study, we suggest that caches improve prey collection and transport in *E. hamatum*. While our hypotheses need to be tested through extensive work in the field, they raise novel questions integrating the growing knowledge of foraging and migration in army ants.

## Methods

We carried out observations in a 220 ha Amazonian primary forest fragment (Terra Firme, coordinates: −1.034113, −46.766017) in the Bragança city area, state of Pará, Brazil. To locate caches, we followed foraging columns across multiple sessions (July 2019-January 2020) between 8:00 am and 4:30 pm, when *E. hamatum* forages [25]. To minimize resampling, we did not collect caches closer than 50 meters. For each cache, we inspected the surrounding 10-meter radius for prey nests or bivouacs, noting whether: it appeared at a multiple-trail junction; it was exposed or covered by leaf litter/fallen tree branches; it was at the side of, or crossed by, trails. We collected all prey from six caches, immediately placing it in 700 ml plastic containers and then storing it sorted by developmental stages (larva, pupa, adult) in 70% ethanol. We later identified prey at the subfamily/genus level using keys for neotropical adult ants [26] and larvae [27], and measured their length.

### Field observations and simulations

We filmed caches for 5 minutes from ~30 cm of height (30 fps, 1920 x 1080px). The frame included individuals arriving from the foraging front and leaving towards the bivouac, as well as individuals passing at the side of caches. We assumed that *E. hamatum* only transported prey from foraging fronts to caches to the bivouac (and not the opposite), and that multiple-prey loads always included only two prey items. We analyzed videos using Boris [28] and developed simulations in Netlogo.

### Statistical analyses

Using STATISTICA v.10, we compared numbers of workers walking in the same direction through CB and FC, considering these as paired data, with Wilcoxon signed-rank tests. For workers walking in both directions, and one- vs. multiple-prey arriving/departing loads, we considered data as unpaired, using Mann Whitney U tests. We analyzed data from simulations using General Linear Models (GLM) in the R [29] package lme4 [30] (Table S7), generating all graphs via ggplot2 [31] and gridExtra [32].

## Supporting information

Data accessibility

Data accessibility

Data accessibility

Data accessibility

Data accessibility

Data accessibility

## Data accessibility

All data are provided in the Supplementary Materials.

## Authors’ contributions

HPDL and RLCDL conceived the study, conducted fieldwork, analyzed fieldwork data and edited the manuscript. NC conceived and supervised the study and wrote the manuscript. ST conceived the study, developed the simulations, analyzed the simulation data and wrote the manuscript. RSFC conceived the study and edited the manuscript. All authors approved the final version of the manuscript and agreed to be held accountable for the content therein.

## Acknowledgments

Marilia Lobão made the study area available for fieldwork. Erika Dawson and Maria Eduarda de Lima Vieira commented on the manuscript. We thank USP and its Experimental Psychology Program, as well as all other universities and postgraduate programs that bravely resist constant attacks on science and scientists in Brazil.

