## Supplementary material for "Temporary prey storage along swarm columns of army ants: an adaptive strategy for successful raiding?": Data accessibility: caches_script_models.R

```
library(lme4)

all_data=read.table(filename <- file.choose(), 
                 header=TRUE,
                 sep=",", 
                 dec=".") # this is the decimal sign, change it to comma if data look weird

# make sure all variables except time and food are considered as factors, visualized in this order in plots

all_data$colsize <- factor(all_data$colsize, levels = c("50","100","300","500","700")) 
all_data$foodevap <- factor(all_data$foodevap, levels = c("0","25","50","75","100")) 
all_data$likeliness <- factor(all_data$likeliness, levels = c("25","50","75","100"))  


### these models compare retrieved prey amounts (caches vs no caches, each treatment group nested in brood evacuation speed and raider number)

glm1 = glm(food ~ foodevap/colsize/treat, subset(all_data, likeliness=="25" & treat != "DOLoff"), family = 'quasipoisson')
summary(glm1)

glm2 = glm(food ~ foodevap/colsize/treat, subset(all_data, likeliness=="50" & treat != "DOLoff"), family = 'quasipoisson')
summary(glm2)

glm3 = glm(food ~ foodevap/colsize/treat, subset(all_data, likeliness=="75" & treat != "DOLoff"), family = 'quasipoisson')
summary(glm3)

glm4 = glm(food ~ foodevap/colsize/treat, subset(all_data, likeliness=="100" & treat != "DOLoff"), family = 'quasipoisson')
summary(glm4)

### these models compare the time spent retrieving prey (caches vs no caches, each treatment group nested in brood evacuation speed and raider number)

tglm1 = glm(time ~ foodevap/colsize/treat, subset(all_data, likeliness=="25" & treat != "DOLoff"), family = 'quasipoisson')
summary(tglm1)

tglm2 = glm(time ~ foodevap/colsize/treat, subset(all_data, likeliness=="50" & treat != "DOLoff"), family = 'quasipoisson')
summary(tglm2)

tglm3 = glm(time ~ foodevap/colsize/treat, subset(all_data, likeliness=="75" & treat != "DOLoff"), family = 'quasipoisson')
summary(tglm3)

tglm4 = glm(time ~ foodevap/colsize/treat, subset(all_data, likeliness=="100" & treat != "DOLoff"), family = 'quasipoisson')
summary(tglm4)

### these models compare the amount of retrieved prey in continuous vs single caching
###(caches vs no caches, each treatment group nested in brood evacuation speed and raider number)

dlglm1 = glm(food ~ foodevap/colsize/treat, subset(all_data, likeliness=="25" & treat != "cachesoff"), family = 'quasipoisson')
summary(dlglm1)

dlglm2 = glm(food ~ foodevap/colsize/treat, subset(all_data, likeliness=="50" & treat != "cachesoff"), family = 'quasipoisson')
summary(dlglm2)

dlglm3 = glm(food ~ foodevap/colsize/treat, subset(all_data, likeliness=="75" & treat != "cachesoff"), family = 'quasipoisson')
summary(dlglm3)

dlglm4 = glm(food ~ foodevap/colsize/treat, subset(all_data, likeliness=="100" & treat != "cachesoff"), family = 'quasipoisson')
summary(dlglm4)

### these models compare time spent retrieving food in continuous vs single caching
###(caches vs no caches, each treatment group nested in brood evacuation speed and raider number)

tdlglm1 = glm(time ~ foodevap/colsize/treat, subset(all_data, likeliness=="25" & treat != "cachesoff"), family = 'quasipoisson')
summary(tdlglm1)

tdlglm2 = glm(time ~ foodevap/colsize/treat, subset(all_data, likeliness=="50" & treat != "cachesoff"), family = 'quasipoisson')
summary(tdlglm2)

tdlglm3 = glm(time ~ foodevap/colsize/treat, subset(all_data, likeliness=="75" & treat != "cachesoff"), family = 'quasipoisson')
summary(tdlglm3)

tdlglm4 = glm(time ~ foodevap/colsize/treat, subset(all_data, likeliness=="100" & treat != "cachesoff"), family = 'quasipoisson')
summary(tdlglm4)


### this model tests effects of brood evacuation speed, raider numbers, probability to cache/collect cached prey items
### response variable is number of retrieved prey items

glmfoodevap = glm(food ~ foodevap + likeliness + colsize, subset(all_data, treat != "DOLoff"), family = 'quasipoisson')
summary(glmfoodevap)


### this model tests effects of brood evacuation speed, raider numbers, probability to cache/collect cached prey items
### response variable is time spent in retrieving all collected prey items

fglmfoodevap = glm(time ~ foodevap + likeliness + colsize, subset(all_data, treat != "DOLoff"), family = 'quasipoisson')
summary(fglmfoodevap)
```
