## Supplementary material for "Temporary prey storage along swarm columns of army ants: an adaptive strategy for successful raiding?": Data accessibility: 2021_07_27_caches_Serafino.nlogo

```
patches-own [
  trail-pheromone       ;; amount of trail-pheromone on a patch, ants release it and go where it is more concentrated, ants carrying prey release more of it
  food?                 ;; true on food patches, false elsewhere
  food2?                ;; true on food patches, false elsewhere
  dropped-food?         ;; true on dropped food patches, false elsewhere
  bivouac?              ;; true/false, it is the green area where bivouac ants stay
]

globals [retrieved-food] ;; this is a counter to keep track of the food arriving at the bivouac, you see it in the plot and as a monitor in the interface

;;;;;;;;;;;;;;;;;;;;;;;;
;;; Setup procedures ;;;
;;;;;;;;;;;;;;;;;;;;;;;;

to setup
  resize-world -50 50 -12 12 ;; makes the world long and narrow
  clear-all
  set-default-shape turtles "bug" ;; gives the agents a 'ant' shape
  create-turtles colony-size ;; creates a number of ants that depends on the corresponding interface slider
  [ set size 2         ;; sets the size of the agents
   set color red       ;; sets the color of the agents
   setxy 0 0  ]        ;; places the origin of the agents, see below
  setup-patches
  reset-ticks
end

to setup-patches  ;; creates the patches corresponding to food etc
  ask patches [ setup-bivouac
    setup-food
    recolor-patch ]
end

to setup-bivouac  ;; patch procedure, bivouac? variable true inside the green area, false elsewhere
  set bivouac? (distancexy 0 0) < 2  ;; positions the nest at left side
end

to setup-food  ;; patch procedure
  set food? (distancexy (0.8 * max-pxcor) 0) < 5  ;; positions the food at right side
  if food? = TRUE [set trail-pheromone trail-pheromone + 1 ]   ;; food has a quantity 1 of trail pheromone (can be thought as smell of prey)
  set food2? (distancexy (-0.8 * max-pxcor) 0) < 5  ;; positions the food at right side
  if food2? = TRUE [set trail-pheromone trail-pheromone + 1 ]   ;; food has a quantity 1 of trail pheromone (can be thought as smell of prey)
end

to recolor-patch  ;; patch procedure, gives color to food
   set pcolor scale-color green trail-pheromone 0.1 5 ;; scale color to show trail-pheromone concentration
  if food? = TRUE [ set pcolor cyan
      ask neighbors [ set trail-pheromone trail-pheromone + 0.1 ]]
  if food2? = TRUE  [ set pcolor sky
  ask neighbors [ set trail-pheromone trail-pheromone + 0.1 ]]
  if dropped-food? = TRUE  [ set pcolor yellow
     set trail-pheromone trail-pheromone + 1
     ask patches in-radius 2 [set trail-pheromone trail-pheromone + 0.1]]   ;;  volatile odor of food on the 8 surronding patches of each food item; this creates a gradient that ants follow, which accelerates food recovery and directs ant lines
  if bivouac? = TRUE [ set pcolor green]
end

to food-disappear ;; this is the food evaporation that simulates prey colonies leaving with their brood
  if ticks mod ((101 - food-evaporation-rate)) = 0
  [ask n-of 1 patches with [ food? = TRUE ]
  [set food? FALSE]]
end

to food2-disappear ;; this is the food evaporation that simulates prey colonies leaving with their brood; still need to organize for the tick thing
  if ticks mod ((101 - food-evaporation-rate)) = 0
  [ask n-of 1 patches with [ food2? = TRUE ]
  [set food2? FALSE]]
end

to count-food ; function that triggers food evaporation when raid starts
  if count patches with [food? = TRUE ] < (70 - individuals-starting-the-raid) and count patches with [food? = TRUE ] > 0 ;; here I have tailored the food patches value on the size of the food patch I use, it can be made proportional if one wants to change food patch size
  [food-disappear]
end

to count-food2 ; function that triggers food evaporation when raid starts
  if count patches with [food2? = TRUE ] < (70 - individuals-starting-the-raid) and count patches with [food2? = TRUE ] > 0 ;; here I have tailored the food patches value on the size of the food patch I use, it can be made proportional if one wants to change food patch size
  [food2-disappear]
end

;;;;;;;;;;;;;;;;;;;;;
;;; Go procedures ;;;
;;;;;;;;;;;;;;;;;;;;;

to go  ;; forever button - starts all the procedures
  if count patches with [food?] + count patches with [food2?] + count patches with [pcolor = yellow] +
  count turtles with [color = pink] + count turtles with [color = orange] + count turtles with [color = violet] = 0 [stop] ;; makes a simulation stop when all food has reached the bivouac
  ask turtles [
   if who >= ticks [ stop ] ;; delay initial departure
   wiggle ;; wiggle side to side while moving
   fd 0.2 ;; advance of 0.2 patches/tick
   set trail-pheromone trail-pheromone + pheromone-released / 100
   if (trail-pheromone >= (101 - pheromone-sensitivity) / 100) [ uphill-trail-pheromone ] ;; each ant follows the trail pheromone towards the higher concentration side; I removed the upper limit here, which incidentally creates loops at low evaporation rates
   if who < colony-size / 100 * percentage-of-bivouac-ants [ bivouac ]   ;; the proportion of ants in the bivouac is selected via a slider from the interface
   ifelse division-of-labor? [ifelse color = red [look-for-food][ifelse color = orange [behave][bring-food-to-bivouac]]] ;; take food where you find many nestmates (orange ants), or home (pink and violet ants)
                             [ifelse color = red [look-for-food][behave]]]  ;; take food where you find many nestmates, or home, pink and violet cache food too
   ask patches [set trail-pheromone trail-pheromone * (100 - pheromone-evaporation-rate) / 100  ;; slowly evaporate trail-pheromone
   recolor-patch]
   count-food
   count-food2 ;;  counts the amount of food
   tick
end

to bivouac ;; makes a portion of ants staying in the green area, these simulate the bivouac
   if [pcolor] of patch-ahead 1 != green [ lt 180 ]
end

to look-for-food  ;; turtle procedure
  if food? = TRUE [set color orange
    set food? FALSE ] ;; pick up food, food disappears drom count, same for food2
   if food2? = TRUE [set color orange
    set food2? FALSE]
   if dropped-food? = TRUE [if random (101 - probability-of-recovering-cached-food ) = 0
    [ set dropped-food? FALSE
      set color pink]]
end

to double-prey
  if color = pink and dropped-food? = TRUE
  [if random (101 - probability-of-recovering-cached-food) = 0 [set color violet
    set dropped-food? FALSE]]
end

to behave ;; makes ants cache food or not, according to the interface switch
ifelse caches? [bring-food-to-nestmates][bring-food-to-bivouac]
end

to bring-food-to-nestmates  ;; turtle procedure; ants deposit food according to density of nestmates, and always when they hit the bivouac
  let density-in-patch count other turtles in-radius 2  ;; creates a local variable counting the number of unloaded/cache collecting ants in the 3px radius of each ant. one could also include orange ants but needs to exclude the raid area then
    if density-in-patch = random ( ifelse-value (density-in-patch = 0)
    [ 1000000 ] ;; if density of nestmates in radius x pixels around an ant = 0, there is one in a million possibility they drop the food they carry
    [100 * (colony-size - density-in-patch) / likeliness-to-cache-food ]) ;; this varies based on colony size
    and food? = FALSE
    and food2? = FALSE
    and pcolor != yellow ;;  so here I originally wanted to code it like 'and dropped-food? = FALSE' but it does not allow ants to cache, no idea why!! but like this it works and it is the same
    or bivouac? = TRUE
    [drop-food]
  double-prey
end

to drop-food ;; turtle procedure ; adds retrieved food to the count if ants hit the bivouac, makes ants cache food otherwise
  ifelse bivouac? = TRUE [ifelse color = violet [set retrieved-food retrieved-food + 2] [set retrieved-food retrieved-food + 1]
    set color red]
  [ set dropped-food? TRUE
    ifelse color != violet [set color red][set color pink]]
end

to bring-food-to-bivouac ;;  turtle procedure ; makes ants drop food in the bivouac, without caches
  if bivouac? [drop-food
    set color red]
    double-prey
end

;; sniff left and right, and go where the strongest smell is
to uphill-trail-pheromone  ;; turtle procedure
  let scent-ahead trail-pheromone-scent-at-angle   0
  let scent-right trail-pheromone-scent-at-angle  45
  let scent-left  trail-pheromone-scent-at-angle -45
  if (scent-right > scent-ahead) or (scent-left > scent-ahead)
  [ ifelse scent-right > scent-left
    [ rt 45 ]
    [ lt 45 ] ]
end

to wiggle  ;; turtle procedure for when ants hit the boarders of the world
  rt random 25
  lt random 25
  if not can-move? 1 [ rt 45] ;; original here was rt 180
end

to-report trail-pheromone-scent-at-angle [angle]
  let p patch-right-and-ahead angle 1
  if p = nobody [ report 0 ]
  report [trail-pheromone] of p
end


; Copyright 1997 Uri Wilensky.  ;; need to check out this
; See Info tab for full copyright and license.
@#$#@#$#@
GRAPHICS-WINDOW
202
300
1187
550
-1
-1
9.674
1
10
1
1
1
0
0
0
1
-50
50
-12
12
1
1
1
ticks
30.0

BUTTON
22
397
102
430
NIL
setup
NIL
1
T
OBSERVER
NIL
NIL
NIL
NIL
1

SLIDER
217
19
445
52
pheromone-evaporation-rate
pheromone-evaporation-rate
10
60
10.0
1.0
1
NIL
HORIZONTAL

BUTTON
112
397
187
430
NIL
go
T
1
T
OBSERVER
NIL
NIL
NIL
NIL
0

SLIDER
20
19
210
52
colony-size
colony-size
0.0
1000.0
100.0
1.0
1
NIL
HORIZONTAL

PLOT
15
99
1352
288
Food
time
food
0.0
200.0
0.0
70.0
true
true
"" ""
PENS
"Available food" 1.0 0 -11221820 true "" "plot count patches with [food? = TRUE]"
"cache recoverers" 1.0 0 -2064490 true "" "plot count turtles with [color = pink]"
"food loaded ants" 1.0 0 -955883 true "" "plot count turtles with [color = orange]"
"dropped food" 1.0 0 -1184463 true "" "plot count patches with [dropped-food? = TRUE]"
"retrieved food" 1.0 0 -7500403 true "" "plot retrieved-food"
"violet ants" 1.0 0 -8630108 true "" "plot count turtles with [color = violet]"
"available food2" 1.0 0 -13791810 true "" "plot count patches with [food2? = TRUE]"

SLIDER
698
15
929
48
percentage-of-bivouac-ants
percentage-of-bivouac-ants
1
100
30.0
1
1
NIL
HORIZONTAL

SLIDER
1099
54
1359
87
probability-of-recovering-cached-food
probability-of-recovering-cached-food
1
100
50.0
1
1
NIL
HORIZONTAL

SLIDER
932
15
1126
48
food-evaporation-rate
food-evaporation-rate
1
100
50.0
1
1
NIL
HORIZONTAL

SLIDER
1132
15
1357
48
individuals-starting-the-raid
individuals-starting-the-raid
1
10
10.0
1
1
NIL
HORIZONTAL

SLIDER
924
54
1095
87
likeliness-to-cache-food
likeliness-to-cache-food
1
100
50.0
1
1
NIL
HORIZONTAL

SWITCH
20
310
125
343
caches?
caches?
1
1
-1000

MONITOR
1244
233
1348
278
NIL
retrieved-food
17
1
11

SLIDER
19
57
213
90
pheromone-sensitivity
pheromone-sensitivity
1
100
30.0
1
1
NIL
HORIZONTAL

SLIDER
220
57
404
90
pheromone-released
pheromone-released
1
100
25.0
1
1
NIL
HORIZONTAL

SWITCH
20
351
190
384
division-of-labor?
division-of-labor?
0
1
-1000

@#$#@#$#@
## WHAT IS IT?

In this project, a colony of ants forages for food. Though each ant follows a set of simple rules, the colony as a whole acts in a sophisticated way.

## HOW IT WORKS

When an ant finds a piece of food, it carries the food back to the nest, dropping a chemical as it moves. When other ants "sniff" the chemical, they follow the chemical toward the food. As more ants carry food to the nest, they reinforce the chemical trail.

## HOW TO USE IT

Click the SETUP button to set up the ant nest (in violet, at center) and three piles of food. Click the GO button to start the simulation. The chemical is shown in a green-to-white gradient.

The EVAPORATION-RATE slider controls the evaporation rate of the chemical. The DIFFUSION-RATE slider controls the diffusion rate of the chemical.

If you want to change the number of ants, move the POPULATION slider before pressing SETUP.

## THINGS TO NOTICE

The ant colony generally exploits the food source in order, starting with the food closest to the nest, and finishing with the food most distant from the nest. It is more difficult for the ants to form a stable trail to the more distant food, since the chemical trail has more time to evaporate and diffuse before being reinforced.

Once the colony finishes collecting the closest food, the chemical trail to that food naturally disappears, freeing up ants to help collect the other food sources. The more distant food sources require a larger "critical number" of ants to form a stable trail.

The consumption of the food is shown in a plot.  The line colors in the plot match the colors of the food piles.

## EXTENDING THE MODEL

Try different placements for the food sources. What happens if two food sources are equidistant from the nest? When that happens in the real world, ant colonies typically exploit one source then the other (not at the same time).

In this project, the ants use a "trick" to find their way back to the nest: they follow the "nest scent." Real ants use a variety of different approaches to find their way back to the nest. Try to implement some alternative strategies.

The ants only respond to chemical levels between 0.05 and 2.  The lower limit is used so the ants aren't infinitely sensitive.  Try removing the upper limit.  What happens?  Why?

In the `uphill-chemical` procedure, the ant "follows the gradient" of the chemical. That is, it "sniffs" in three directions, then turns in the direction where the chemical is strongest. You might want to try variants of the `uphill-chemical` procedure, changing the number and placement of "ant sniffs."

## NETLOGO FEATURES

The built-in `diffuse` primitive lets us diffuse the chemical easily without complicated code.

The primitive `patch-right-and-ahead` is used to make the ants smell in different directions without actually turning.

## HOW TO CITE

If you mention this model or the NetLogo software in a publication, we ask that you include the citations below.

For the model itself:

* Wilensky, U. (1997).  NetLogo Ants model.  http://ccl.northwestern.edu/netlogo/models/Ants.  Center for Connected Learning and Computer-Based Modeling, Northwestern University, Evanston, IL.

Please cite the NetLogo software as:

* Wilensky, U. (1999). NetLogo. http://ccl.northwestern.edu/netlogo/. Center for Connected Learning and Computer-Based Modeling, Northwestern University, Evanston, IL.

## COPYRIGHT AND LICENSE

Copyright 1997 Uri Wilensky.

![CC BY-NC-SA 3.0](http://ccl.northwestern.edu/images/creativecommons/byncsa.png)

This work is licensed under the Creative Commons Attribution-NonCommercial-ShareAlike 3.0 License.  To view a copy of this license, visit https://creativecommons.org/licenses/by-nc-sa/3.0/ or send a letter to Creative Commons, 559 Nathan Abbott Way, Stanford, California 94305, USA.

Commercial licenses are also available. To inquire about commercial licenses, please contact Uri Wilensky at.

This model was created as part of the project: CONNECTED MATHEMATICS: MAKING SENSE OF COMPLEX PHENOMENA THROUGH BUILDING OBJECT-BASED PARALLEL MODELS (OBPML).  The project gratefully acknowledges the support of the National Science Foundation (Applications of Advanced Technologies Program) -- grant numbers RED #9552950 and REC #9632612.

This model was developed at the MIT Media Lab using CM StarLogo.  See Resnick, M. (1994) "Turtles, Termites and Traffic Jams: Explorations in Massively Parallel Microworlds."  Cambridge, MA: MIT Press.  Adapted to StarLogoT, 1997, as part of the Connected Mathematics Project.

This model was converted to NetLogo as part of the projects: PARTICIPATORY SIMULATIONS: NETWORK-BASED DESIGN FOR SYSTEMS LEARNING IN CLASSROOMS and/or INTEGRATED SIMULATION AND MODELING ENVIRONMENT. The project gratefully acknowledges the support of the National Science Foundation (REPP & ROLE programs) -- grant numbers REC #9814682 and REC-0126227. Converted from StarLogoT to NetLogo, 1998.

<!-- 1997 1998 MIT -->
@#$#@#$#@
default
true
0
Polygon -7500403 true true 150 5 40 250 150 205 260 250

airplane
true
0
Polygon -7500403 true true 150 0 135 15 120 60 120 105 15 165 15 195 120 180 135 240 105 270 120 285 150 270 180 285 210 270 165 240 180 180 285 195 285 165 180 105 180 60 165 15

arrow
true
0
Polygon -7500403 true true 150 0 0 150 105 150 105 293 195 293 195 150 300 150

box
false
0
Polygon -7500403 true true 150 285 285 225 285 75 150 135
Polygon -7500403 true true 150 135 15 75 150 15 285 75
Polygon -7500403 true true 15 75 15 225 150 285 150 135
Line -16777216 false 150 285 150 135
Line -16777216 false 150 135 15 75
Line -16777216 false 150 135 285 75

bug
true
0
Circle -7500403 true true 96 182 108
Circle -7500403 true true 110 127 80
Circle -7500403 true true 110 75 80
Line -7500403 true 150 100 80 30
Line -7500403 true 150 100 220 30

butterfly
true
0
Polygon -7500403 true true 150 165 209 199 225 225 225 255 195 270 165 255 150 240
Polygon -7500403 true true 150 165 89 198 75 225 75 255 105 270 135 255 150 240
Polygon -7500403 true true 139 148 100 105 55 90 25 90 10 105 10 135 25 180 40 195 85 194 139 163
Polygon -7500403 true true 162 150 200 105 245 90 275 90 290 105 290 135 275 180 260 195 215 195 162 165
Polygon -16777216 true false 150 255 135 225 120 150 135 120 150 105 165 120 180 150 165 225
Circle -16777216 true false 135 90 30
Line -16777216 false 150 105 195 60
Line -16777216 false 150 105 105 60

car
false
0
Polygon -7500403 true true 300 180 279 164 261 144 240 135 226 132 213 106 203 84 185 63 159 50 135 50 75 60 0 150 0 165 0 225 300 225 300 180
Circle -16777216 true false 180 180 90
Circle -16777216 true false 30 180 90
Polygon -16777216 true false 162 80 132 78 134 135 209 135 194 105 189 96 180 89
Circle -7500403 true true 47 195 58
Circle -7500403 true true 195 195 58

circle
false
0
Circle -7500403 true true 0 0 300

circle 2
false
0
Circle -7500403 true true 0 0 300
Circle -16777216 true false 30 30 240

cow
false
0
Polygon -7500403 true true 200 193 197 249 179 249 177 196 166 187 140 189 93 191 78 179 72 211 49 209 48 181 37 149 25 120 25 89 45 72 103 84 179 75 198 76 252 64 272 81 293 103 285 121 255 121 242 118 224 167
Polygon -7500403 true true 73 210 86 251 62 249 48 208
Polygon -7500403 true true 25 114 16 195 9 204 23 213 25 200 39 123

cylinder
false
0
Circle -7500403 true true 0 0 300

dot
false
0
Circle -7500403 true true 90 90 120

face happy
false
0
Circle -7500403 true true 8 8 285
Circle -16777216 true false 60 75 60
Circle -16777216 true false 180 75 60
Polygon -16777216 true false 150 255 90 239 62 213 47 191 67 179 90 203 109 218 150 225 192 218 210 203 227 181 251 194 236 217 212 240

face neutral
false
0
Circle -7500403 true true 8 7 285
Circle -16777216 true false 60 75 60
Circle -16777216 true false 180 75 60
Rectangle -16777216 true false 60 195 240 225

face sad
false
0
Circle -7500403 true true 8 8 285
Circle -16777216 true false 60 75 60
Circle -16777216 true false 180 75 60
Polygon -16777216 true false 150 168 90 184 62 210 47 232 67 244 90 220 109 205 150 198 192 205 210 220 227 242 251 229 236 206 212 183

fish
false
0
Polygon -1 true false 44 131 21 87 15 86 0 120 15 150 0 180 13 214 20 212 45 166
Polygon -1 true false 135 195 119 235 95 218 76 210 46 204 60 165
Polygon -1 true false 75 45 83 77 71 103 86 114 166 78 135 60
Polygon -7500403 true true 30 136 151 77 226 81 280 119 292 146 292 160 287 170 270 195 195 210 151 212 30 166
Circle -16777216 true false 215 106 30

flag
false
0
Rectangle -7500403 true true 60 15 75 300
Polygon -7500403 true true 90 150 270 90 90 30
Line -7500403 true 75 135 90 135
Line -7500403 true 75 45 90 45

flower
false
0
Polygon -10899396 true false 135 120 165 165 180 210 180 240 150 300 165 300 195 240 195 195 165 135
Circle -7500403 true true 85 132 38
Circle -7500403 true true 130 147 38
Circle -7500403 true true 192 85 38
Circle -7500403 true true 85 40 38
Circle -7500403 true true 177 40 38
Circle -7500403 true true 177 132 38
Circle -7500403 true true 70 85 38
Circle -7500403 true true 130 25 38
Circle -7500403 true true 96 51 108
Circle -16777216 true false 113 68 74
Polygon -10899396 true false 189 233 219 188 249 173 279 188 234 218
Polygon -10899396 true false 180 255 150 210 105 210 75 240 135 240

house
false
0
Rectangle -7500403 true true 45 120 255 285
Rectangle -16777216 true false 120 210 180 285
Polygon -7500403 true true 15 120 150 15 285 120
Line -16777216 false 30 120 270 120

leaf
false
0
Polygon -7500403 true true 150 210 135 195 120 210 60 210 30 195 60 180 60 165 15 135 30 120 15 105 40 104 45 90 60 90 90 105 105 120 120 120 105 60 120 60 135 30 150 15 165 30 180 60 195 60 180 120 195 120 210 105 240 90 255 90 263 104 285 105 270 120 285 135 240 165 240 180 270 195 240 210 180 210 165 195
Polygon -7500403 true true 135 195 135 240 120 255 105 255 105 285 135 285 165 240 165 195

line
true
0
Line -7500403 true 150 0 150 300

line half
true
0
Line -7500403 true 150 0 150 150

pentagon
false
0
Polygon -7500403 true true 150 15 15 120 60 285 240 285 285 120

person
false
0
Circle -7500403 true true 110 5 80
Polygon -7500403 true true 105 90 120 195 90 285 105 300 135 300 150 225 165 300 195 300 210 285 180 195 195 90
Rectangle -7500403 true true 127 79 172 94
Polygon -7500403 true true 195 90 240 150 225 180 165 105
Polygon -7500403 true true 105 90 60 150 75 180 135 105

plant
false
0
Rectangle -7500403 true true 135 90 165 300
Polygon -7500403 true true 135 255 90 210 45 195 75 255 135 285
Polygon -7500403 true true 165 255 210 210 255 195 225 255 165 285
Polygon -7500403 true true 135 180 90 135 45 120 75 180 135 210
Polygon -7500403 true true 165 180 165 210 225 180 255 120 210 135
Polygon -7500403 true true 135 105 90 60 45 45 75 105 135 135
Polygon -7500403 true true 165 105 165 135 225 105 255 45 210 60
Polygon -7500403 true true 135 90 120 45 150 15 180 45 165 90

square
false
0
Rectangle -7500403 true true 30 30 270 270

square 2
false
0
Rectangle -7500403 true true 30 30 270 270
Rectangle -16777216 true false 60 60 240 240

star
false
0
Polygon -7500403 true true 151 1 185 108 298 108 207 175 242 282 151 216 59 282 94 175 3 108 116 108

target
false
0
Circle -7500403 true true 0 0 300
Circle -16777216 true false 30 30 240
Circle -7500403 true true 60 60 180
Circle -16777216 true false 90 90 120
Circle -7500403 true true 120 120 60

tree
false
0
Circle -7500403 true true 118 3 94
Rectangle -6459832 true false 120 195 180 300
Circle -7500403 true true 65 21 108
Circle -7500403 true true 116 41 127
Circle -7500403 true true 45 90 120
Circle -7500403 true true 104 74 152

triangle
false
0
Polygon -7500403 true true 150 30 15 255 285 255

triangle 2
false
0
Polygon -7500403 true true 150 30 15 255 285 255
Polygon -16777216 true false 151 99 225 223 75 224

truck
false
0
Rectangle -7500403 true true 4 45 195 187
Polygon -7500403 true true 296 193 296 150 259 134 244 104 208 104 207 194
Rectangle -1 true false 195 60 195 105
Polygon -16777216 true false 238 112 252 141 219 141 218 112
Circle -16777216 true false 234 174 42
Rectangle -7500403 true true 181 185 214 194
Circle -16777216 true false 144 174 42
Circle -16777216 true false 24 174 42
Circle -7500403 false true 24 174 42
Circle -7500403 false true 144 174 42
Circle -7500403 false true 234 174 42

turtle
true
0
Polygon -10899396 true false 215 204 240 233 246 254 228 266 215 252 193 210
Polygon -10899396 true false 195 90 225 75 245 75 260 89 269 108 261 124 240 105 225 105 210 105
Polygon -10899396 true false 105 90 75 75 55 75 40 89 31 108 39 124 60 105 75 105 90 105
Polygon -10899396 true false 132 85 134 64 107 51 108 17 150 2 192 18 192 52 169 65 172 87
Polygon -10899396 true false 85 204 60 233 54 254 72 266 85 252 107 210
Polygon -7500403 true true 119 75 179 75 209 101 224 135 220 225 175 261 128 261 81 224 74 135 88 99

wheel
false
0
Circle -7500403 true true 3 3 294
Circle -16777216 true false 30 30 240
Line -7500403 true 150 285 150 15
Line -7500403 true 15 150 285 150
Circle -7500403 true true 120 120 60
Line -7500403 true 216 40 79 269
Line -7500403 true 40 84 269 221
Line -7500403 true 40 216 269 79
Line -7500403 true 84 40 221 269

x
false
0
Polygon -7500403 true true 270 75 225 30 30 225 75 270
Polygon -7500403 true true 30 75 75 30 270 225 225 270
@#$#@#$#@
NetLogo 6.2.0
@#$#@#$#@
@#$#@#$#@
@#$#@#$#@
<experiments>
  <experiment name="experiment" repetitions="25" runMetricsEveryStep="true">
    <setup>setup</setup>
    <go>repeat 500 [ go ]</go>
    <metric>count turtles with [color = pink]</metric>
    <metric>count turtles with [color = orange]</metric>
    <metric>count patches with [pcolor = yellow]</metric>
    <metric>count patches with [pcolor = sky]</metric>
    <metric>count patches with [pcolor = cyan]</metric>
    <metric>retrieved-food</metric>
    <enumeratedValueSet variable="division-of-labor?">
      <value value="true"/>
    </enumeratedValueSet>
    <enumeratedValueSet variable="likeliness-to-cache-food">
      <value value="100"/>
    </enumeratedValueSet>
    <enumeratedValueSet variable="probability-of-recovering-cached-food">
      <value value="100"/>
    </enumeratedValueSet>
    <enumeratedValueSet variable="pheromone-released">
      <value value="25"/>
    </enumeratedValueSet>
    <enumeratedValueSet variable="food-evaporation-rate">
      <value value="50"/>
    </enumeratedValueSet>
    <enumeratedValueSet variable="individuals-starting-the-raid">
      <value value="10"/>
    </enumeratedValueSet>
    <enumeratedValueSet variable="pheromone-evaporation-rate">
      <value value="10"/>
    </enumeratedValueSet>
    <enumeratedValueSet variable="colony-size">
      <value value="50"/>
      <value value="100"/>
    </enumeratedValueSet>
    <enumeratedValueSet variable="caches?">
      <value value="true"/>
      <value value="false"/>
    </enumeratedValueSet>
    <enumeratedValueSet variable="percentage-of-bivouac-ants">
      <value value="30"/>
    </enumeratedValueSet>
    <enumeratedValueSet variable="pheromone-sensitivity">
      <value value="30"/>
    </enumeratedValueSet>
  </experiment>
</experiments>
@#$#@#$#@
@#$#@#$#@
default
0.0
-0.2 0 0.0 1.0
0.0 1 1.0 0.0
0.2 0 0.0 1.0
link direction
true
0
Line -7500403 true 150 150 90 180
Line -7500403 true 150 150 210 180
@#$#@#$#@
0
@#$#@#$#@
```
