## Supplementary figures and images for "Temporary prey storage along swarm columns of army ants: an adaptive strategy for successful raiding?"

### Figure S1

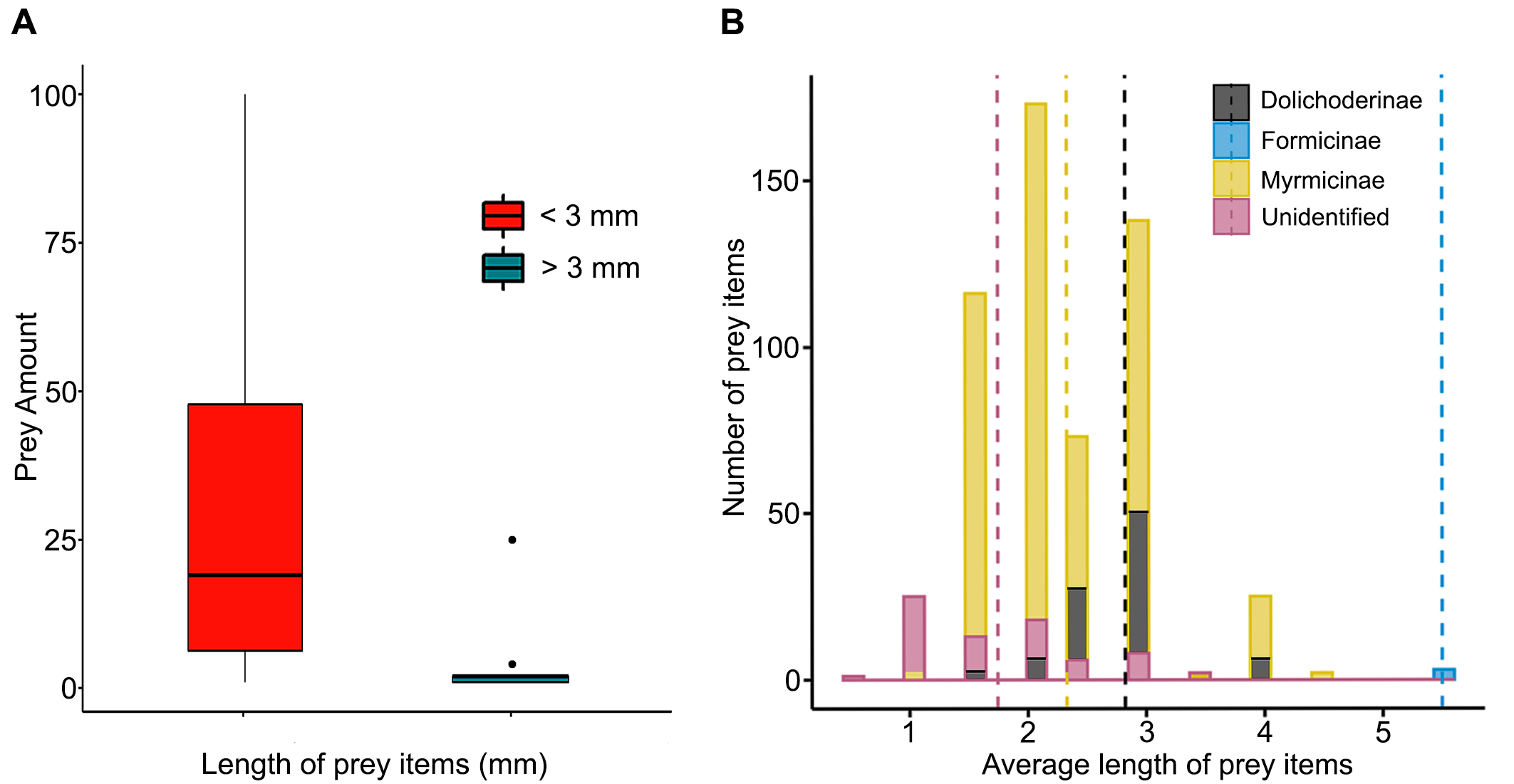

### Figure S2

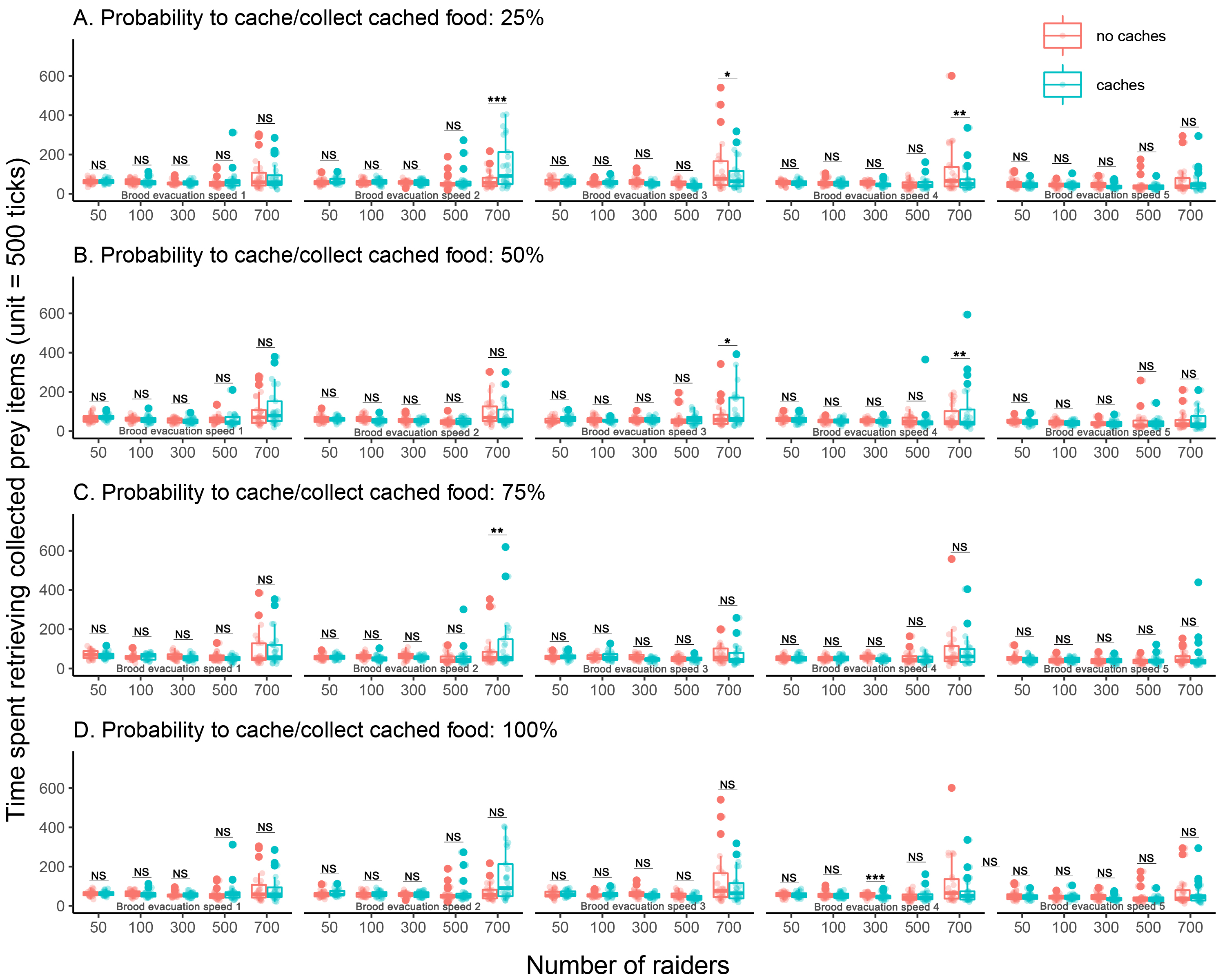

### Figure S3

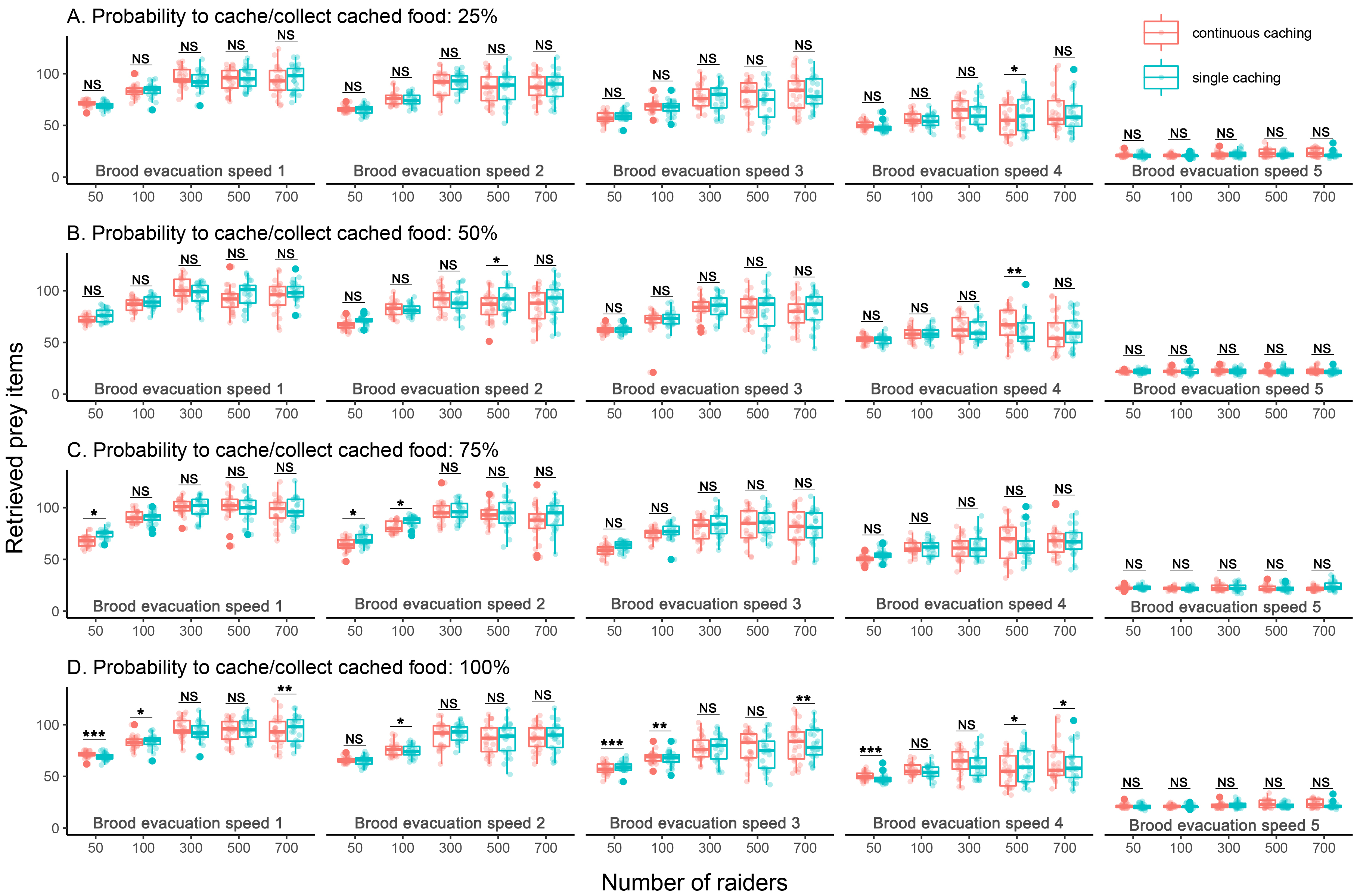

### Figure S4

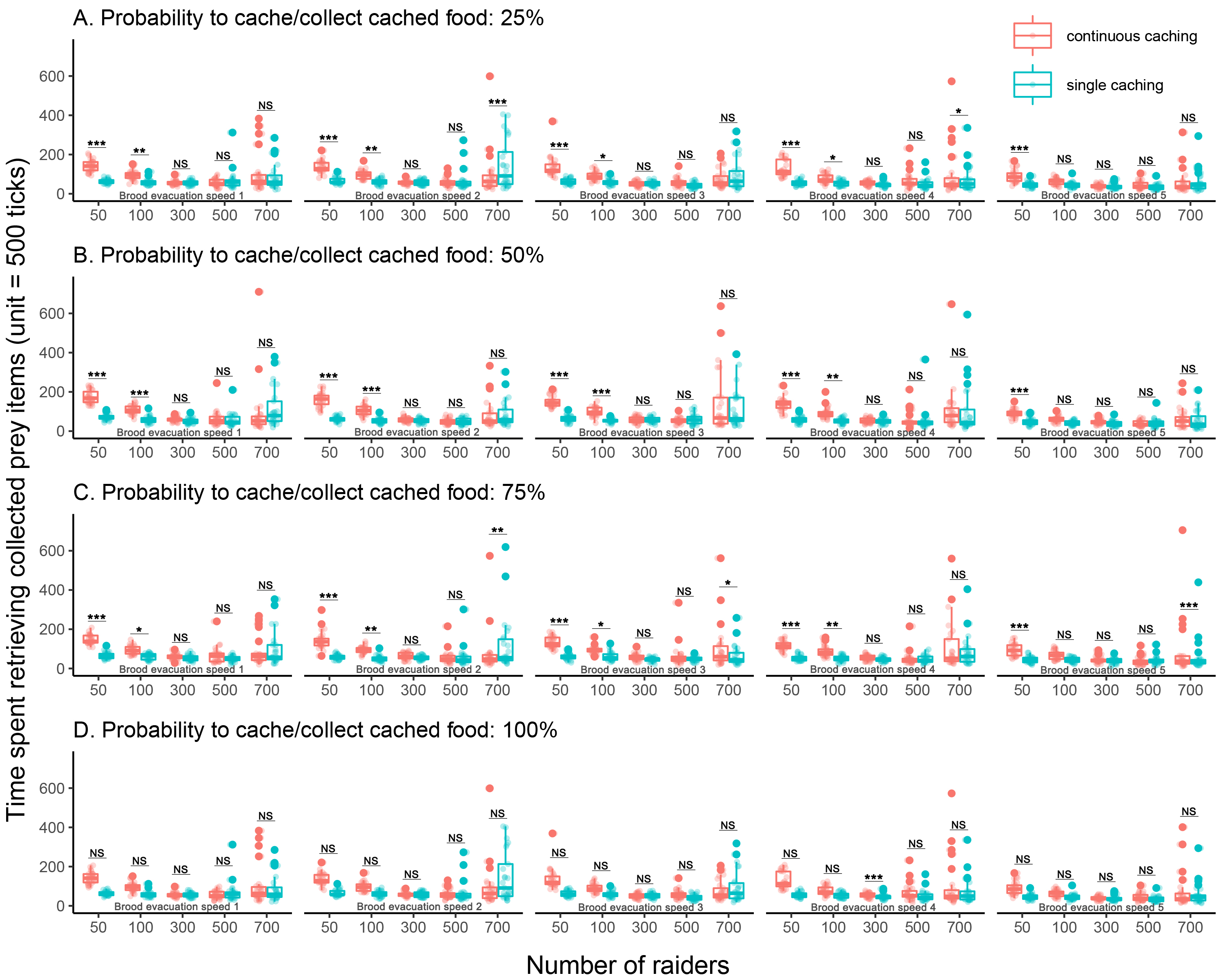
