## Supplementary material for "Temporary prey storage along swarm columns of army ants: an adaptive strategy for successful raiding?": Data accessibility

**Cache characterization and prey diversity.** Of the ten caches we found, five were located at the side of foraging trails and the other five were crossed by foraging trails (Table S1). Four were partially exposed (we could see workers manipulating prey or, in a single case, feeding on it), while six were hidden in the leaf litter or under logs. Nine were in potential bottleneck locations (three after trail junctions and six near logs, fallen branches or roots obstructing the trails). After collecting all prey from 6 caches, we identified 624 specimens (277 larvae, 334 pupae, 86 adults of which 42 males and 5 gynes) as from the ant subfamilies Myrmicinae (530), Dolichoderinae (91) and Formicinae (3). Among those, we assigned 362 specimens to the genera *Pheidole* (304), *Linepithema* (55) and *Camponotus* (3). Substantial damage to prey items (probably caused by *E. hamatum* workers) prevented us from identifying 73 specimens. Prey size ranged from 0.5 to 5.5 mm (2.34±0.74 mm), with gynes of *Camponotus* (3) and Myrmicinae (2) as the largest specimens (respectively, 5.5 and 4 mm) (Figure S2E). Interestingly, two caches included prey of multiple ant subfamilies (Formicinae and Myrmicinae; Dolichoderinae, Formicinae and Myrmicinae, respectively), indicating that single caches include prey collected across multiple raids.

**Simulations.** We used Netlogo [[1]](https://paperpile.com/c/3aC6Th/bJGt1) to develop agent-based simulations (Video S2, model specifications in Table S3, Netlogo model file provided). In each simulation, prey-unloaded ants (body color: red) emerged from the central point (simulating a trail bifurcation or the bivouac itself) of a 101x25-patch 2D world, and walked randomly at a speed of 0.2 patches/time unit (tick). When ants hit the border of the world, they changed direction at a random angle between 1 and 45 degrees. Ants simultaneously followed and deposited a volatile trail pheromone evaporating with time (pheromone code modified from Wilensky 1997 [[2]](https://paperpile.com/c/3aC6Th/qHih)), which resulted in the formation of relatively stable columns of ants walking in both directions. Similar to what happens in real army ants, each virtual ant at the column’s distal end generally re-entered the column guided by the pheromone gradient, after walking a small distance outside of the column itself. This resulted in the advancement of the whole column. A proportion of ants was confined in a 5x5-patch area around the central point of the world (highlighted in green), which resulted in ants being generally attracted by the high concentration of individuals constantly depositing the trail pheromone. In turn, this made ants generally transport prey to the central area and ruled out the need of explicitly modelling a homing behavior. Each of two roughly circular “food piles” situated at the opposite sides of the world included 69 patches (each representing a prey item). Prey items, corresponding to sky- or cyan-colored patches, released an attractive pheromone in the surrounding 1-patch radius, simulating a volatile prey odor. When red ants carrying no prey items walked on a prey item patch, they always collected it, becoming orange. Ants then deposited the collected prey items with a probability increasing with the density of ants they encountered within a 2-patch radius, modulated via a custom parameter (‘likeliness-to-cache-food’) modifiable from the Netlogo interface.

Cached prey items corresponded to yellow patches and released an attractive pheromone on the surrounding 1-patch radius, again simulating a volatile prey odor. Red ants carrying no prey items collected encountered cached food items with a manually regulated probability (interface slider ‘probability-to-collect-cached-food’), becoming pink; pink ants, which always carried a previously cached item, could collect a second prey item with the same probability, becoming violet. As ants collected a threshold number of prey items (always 10 in our simulations, but customizable from the Netlogo interface via the ‘individuals-starting-the-raid’ slider) from a prey pile, indicating the initiation of a raid, random uncollected prey items started disappearing at arbitrary intervals (in our simulation, a random prey item per 100, 75, 50, 25, 1 time units, corresponding to ‘Brood evacuation speed’ 1 to 5, Figure 2). Prey item disappearance simulated the evacuation of brood (the army ant prey) by an attacked ant colony. As the indicator of a raid’s initiation, we used a threshold value (10) of collected prey items, and not a single prey item. We did so because, in our preliminary observations, single unloaded ants occasionally set free from the trail columns and reached the prey piles, triggering the disappearance of prey without a party of ants initiating a raid. In other words, we used such a threshold value to avoid brood evacuation to be triggered by a single or a few ‘scout’ foragers. As a consequence of this threshold value, the minimum amount of retrieved prey items in our simulation was always 20, i.e., 10 for each of two prey piles.

Orange ants carrying 1 prey item, pink ants carrying 1 previously cached item, and violet ants carrying 2 previously cached items added respectively 1, 1, and 2 units to the total count of retrieved prey as they reached the world’s central safe green area. When they did so, they reverted to the red color and were again able to collect prey or cached prey items. If we switched off the caching behavior from a custom command in the Netlogo interface (caches?), orange ants never cached their prey items, and prey retrieval only occurred when they deposited single prey items in the green central area of the world. A second switch, which we named ‘Division-of-labor?’ determined whether collectors of one or two cached prey items (pink or violet ants) kept caching their loads based on the surrounding ant density (Division-of-labor?: OFF), or only when they reached the central green zone, with only orange raiders caching their loads (Division-of-labor?: ON). We refer to these two scenarios as ‘continuous caching’ and ‘single caching’, respectively, in the Supplementary Figures and Tables.

Although parameters can be arbitrarily set via a series of sliders within the Netlogo interface, we decided to conduct our simulations with a limited combination of settings (Table S3). We did so with the aim of reproducing the conditions that best simulated real army ants raids, e.g., stable but not extremely strict columns, and to simplify the interpretation of results. For example, we only used equal probabilities of caching and collecting cached prey items (e.g., probability of caching a food item: 50%; probability to collect a cached food item: 50%), which limited the number of setting combinations. We conducted our simulations using the Netlogo-integrated software Behaviorspace, implementing sessions of multiple simulations over a variety of predefined settings. Considering all parameter combinations pooled, we conducted a total of 7500 simulations (2500 for caching raiders with single caching; 2500 for caching raiders with continuous caching; 2500 for non-caching raiders). We then edited and compiled the Behaviorspace output .csv files via a custom R script, producing a single data sheet (provided) including information about raid size, probability to cache/collect cached food, brood evacuation speed, number of prey items retrieved and time spent to retrieve all collected prey items. To analyze data from simulations, we used Generalized Linear Models (GLM) with Quasipoisson distribution via in the R [[3]](https://paperpile.com/c/3aC6Th/cZlaE) package lme4 [[4]](https://paperpile.com/c/3aC6Th/sbmZb), after assessing that the models’ residual deviance/residual degrees of freedom ratio was higher than ~1, correcting for overdispersion. Response variables and dependent variables for each model are provided in table S7.

**Figures**

**Figure S1. A.** Number of cached prey items by length classes (< 3 mm and > 3mm, n=697). In whisker plots, central lines, boxes, whiskers and dots respectively show median, quartiles, max/min and outliers. **B.** Length distribution of prey items in collected caches by subfamily. Dotted lines represent the mean value for each subfamily. In whisker plots, central lines, boxes, whiskers and dots respectively show median, quartiles, max/min and outliers. ***: p < 0.001, **: p < 0.01, *: p < 0.05; NS: no significance.

**Figure S2.** Simulation results. Time caching vs. non-caching virtual ants spent retrieving collected prey items. In whisker plots (each representing 25 simulations), central lines, boxes, whiskers and dots respectively show median, quartiles, max/min and outliers. ***: p < 0.001, **: p < 0.01, *: p < 0.05; NS: no significance.

**Figure S3.** Simulation results. Prey items retrieved by virtual ants caching food a single time vs. caching food continuously. In whisker plots (each representing 25 simulations), central lines, boxes, whiskers and dots respectively show median, quartiles, max/min and outliers. ***: p < 0.001, **: p < 0.01, *: p < 0.05; NS: no significance.

**Figure S4.** Simulation results. Time spent retrieving collected prey items. Virtual ants caching food a single time vs. caching food continuously. In whisker plots (each representing 25 simulations), central lines, boxes, whiskers and dots respectively show median, quartiles, max/min and outliers. ***: p < 0.001, **: p < 0.01, *: p < 0.05; NS: no significance.
