## Supplementary material for "Temporary prey storage along swarm columns of army ants: an adaptive strategy for successful raiding?": Table S1

**Table S1.** *Eciton hamatum* cache characteristics (leaf litter in primary rainforest floor, Bragança, Pará, Brazil)

| Cache ID | Cache Date | Cache Time | Cache Position | Cache Area | Cache Situation |
|---|---|---|---|---|---|
| Cache 1 | 01/07/2020 | 11h04 am | Side of the trail | Near a Trunk, Branch or Root | Sheltered |
| Cache 2 | 07/09/2019 | 10h48 am | Side of the trail | Near a Trunk, Branch or Root | Sheltered |
| Cache 3 | 07/09/2019 | 10h35 am | Side of the trail | Near a Trunk, Branch or Root | Sheltered |
| Cache 4 | 01/07/2020 | 10h56 am | Side of the trail | Near a Trunk, Branch or Root | Sheltered |
| Cache 5 | 12/17/2019 | 02h41 pm | Side of the trail | Two trails merge | Sheltered |
| Cache 6 | 12/17/2019 | 04h17 pm | Crossed by foraging trails | Two trails merge | Partially Exposed |
| Cache 7 | 12/20/2019 | 02h52 pm | Crossed by foraging trails | Near a Trunk, Branch or Root | Partially Exposed |
| Cache 8 | 10/22/2019 | 09h10 am | Crossed by foraging trails | No obstacles or trail merge visible | Partially Exposed |
| Cache 9 | 10/22/2019 | 09h53 am | Crossed by foraging trails | Two trails merge | Partially Exposed |
| Cache 10 | 10/31/2019 | 04h47 pm | Crossed by foraging trails | Near a Trunk, Branch or Root | Sheltered |
