## Supplementary material for "Temporary prey storage along swarm columns of army ants: an adaptive strategy for successful raiding?": Table S2

**Table S2 -** Collected Cache Prey Size and Video Cache Registration Before and After Cache

| | Cache 1 | Cache 2 | Cache 3 | Cache 4 | Cache 5 | Cache 6 | Cache 7 | Cache 8 | Cache 9 | Cache 10 | Cache 11 | Cache 12 |
|---|---|---|---|---|---|---|---|---|---|---|---|---|
| **Region Before Cache - One Prey** | **n/a** | **n/a** | **24** | **6** | **11** | **17** | **197** | **128** | **0** | **45** | **64** | **80** |
| **Region After Cache - One Prey** | **n/a** | **n/a** | **8** | **2** | **7** | **3** | **125** | **149** | **1** | **25** | **8** | **28** |
| **Region Before Cache - Multiple preys** | **n/a** | **n/a** | **15** | **9** | **3** | **21** | **17** | **21** | **0** | **0** | **2** | **1** |
| **Region After Cache - Multiple preys** | **n/a** | **n/a** | **9** | **4** | **5** | **3** | **30** | **25** | **0** | **0** | **0** | **0** |
| **Region Before Cache - Without Prey** | **n/a** | **n/a** | **123** | **154** | **584** | **71** | **50** | **102** | **28** | **55** | **133** | **214** |
| **Region After Cache - Without Prey** | **n/a** | **n/a** | **68** | **94** | **425** | **47** | **52** | **28** | **49** | **109** | **75** | **218** |
| **Collected Cache - Prey Size: 0,5 mm** | **0** | **0** | **0** | **0** | **1** | **0** | **n/a** | **n/a** | **n/a** | **n/a** | **n/a** | **n/a** |
| **Collected Cache - Prey Size: 1 mm** | **0** | **1** | **0** | **0** | **8** | **18** | **n/a** | **n/a** | **n/a** | **n/a** | **n/a** | **n/a** |
| **Collected Cache - Prey Size: - 1,5 mm** | **38** | **2** | **0** | **0** | **45** | **46** | **n/a** | **n/a** | **n/a** | **n/a** | **n/a** | **n/a** |
| **Collected Cache - Prey Size: 2 mm** | **0** | **4** | **0** | **0** | **100** | **93** | **n/a** | **n/a** | **n/a** | **n/a** | **n/a** | **n/a** |
| **Collected Cache - Prey Size: 2,5 mm** | **0** | **20** | **0** | **7** | **62** | **17** | **n/a** | **n/a** | **n/a** | **n/a** | **n/a** | **n/a** |
| **Collected Cache - Prey Size: 3 mm** | **1** | **0** | **41** | **9** | **53** | **92** | **n/a** | **n/a** | **n/a** | **n/a** | **n/a** | **n/a** |
| **Collected Cache - Prey Size: 3,5 mm** | **0** | **0** | **0** | **1** | **0** | **2** | **n/a** | **n/a** | **n/a** | **n/a** | **n/a** | **n/a** |
| **Collected Cache - Prey Size: 4 mm** | **0** | **0** | **4** | **1** | **1** | **25** | **n/a** | **n/a** | **n/a** | **n/a** | **n/a** | **n/a** |
| **Collected Cache - Prey Size: 4,5 mm** | **0** | **0** | **0** | **0** | **0** | **2** | **n/a** | **n/a** | **n/a** | **n/a** | **n/a** | **n/a** |
| **Collected Cache - Prey Size: 5,5 mm** | **0** | **0** | **0** | **0** | **2** | **1** | **n/a** | **n/a** | **n/a** | **n/a** | **n/a** | **n/a** |
