## Supplementary material for "Temporary prey storage along swarm columns of army ants: an adaptive strategy for successful raiding?": Table S3

| Variable | Values/type | Description |
|---|---|---|
| **Agents** | | |
| X and Y coordinates | Non-integer; X coordinates: -50 – 50 Y coordinates: −12 – 12 | all agents have explicit locations |
| State | "looking for food"; "bringing food to nestmates"or "bringing food to bivouac", both nested in "behave" | ants follow and release a trail pheromone at all times |
| **Patches** | | |
| X and Y coordinates | Non-integer; X coordinates: -50 – 50 Y coordinates: −12 – 12 | every patch has an explicit location and cannot move |
| Patch type | "Nothing", "food", "food2", "bivouac", "dropped-food" | "nothing" defines empty environment; "food1" and "food2" are piled prey items; "bivouac" is the area where a percentage of resident ants is confined at all times, marked with patch color green; "dropped-food" represent cached prey items |
| Trail pheromone | integer, ≥0 | The amount of trail pheromone on a patch |
| individuals-starting-the-raid | integer,1-10 (modelled as 10) | number of collected food or food2 patches determining the stat of a raid and triggering the evacuation of uncollected prey items |
| **Global variables** | | |
| colony-size | integer,1-1000 (modelled as 50, 100, 300, 500, 700) | number of raiding individuals, including those resident in the central green area |
| pheromone-evaporation rate | integer, 10-60 (modelled as 10) | evaporation of the trail pheromone through time |
| pheromone-sensitivity | integer,1-100 (modelled as 30) | level of attraction to the trail pheromone |
| pheromone-released | integer,1-100 (modelled as 25) | quantity of trail pheromone released by each ant |
| percentage-of-bivouac-ants | integer,1-100 (modelled as 30) | percentage of individuals confined in the green central area |
| food-evaporation-rate | integer,1-100 (modelled as 1, 25, 50, 75, 100) | called brood evacuation speed in the text. It is the number of ticks passing between two uncollected prey item disappearance events |
| likeliness-to-cache-food | integer,1-100 (modelled as 25, 50, 75, 100) | probability to cache food according to the density of surrounding ants |
| probability-of-recovering-cached-food | integer,1-100 (modelled as 25, 50, 75, 100) | probability to collect a cached food item when stepping on the corresponding "dropped-food" yellow patch |
| caches | boolean (ON-OFF) | determines whether raiders cache or not |
| division-of-labor | boolean (ON-OFF) | determines whether cache retrievers keep caching based on ant density or only drop prey items on the central green area |
| retrieved-food | integer, ≥0 | the amount of prey items reaching the green area at the center of the grid |

| Variable | Values/type | Description |
|---|---|---|
| **Procedures** | | |
| setup | observer procedure | sets up the simulation world |
| setup-patches | observer procedure | sets up patches |
| setup-bivouac | patch procedure | sets up the green area at the world's center |
| setup-food | patch procedure | sets up prey item piles |
| recolor-patch | patch procedure | maintains/changes color of different types of patches |
| food-disappear | observer procedure | evacuates prey items in pile 1 |
| food2-disappear | observer procedure | evacuates prey items in pile 2 |
| count-food | observer procedure | counts prey items in pile 1 |
| count-food2 | observer procedure | counts prey items in pile 2 |
| go | observer procedure | starts and keeps the simulation going |
| bivouac | turtle procedure | confines a proportion of ants in the central green area |
| look-for-food | turtle procedure | makes red ants move randomly in the world, becoming orange if they step on a food/food2 patch or pink if they encounter a dropped-food patch (with probability = probability-of-recovering-cached-food) |
| double-prey | turtle procedure | makes pink cache-collecting ants collect a second cached item, becoming violet |
| behave | turtle procedure | determines whether ants cache or not |
| bring-food-to-nestmates | turtle procedure | makes ants cache food based on ant density in the surrounding patches and on a fixed probability to cache |
| drop-food | turtle procedure | makes ants drop food and become either red if they are pink or orange, or pink if they are violet. If executing agents are on the central green area, 1 or 2 units are added to the count of the retrieved food |
| bring-food-to-bivouac | turtle procedure | makes ants drop their loads exclusively in the central green area |
| uphill-trail-pheromone | turtle procedure | makes ants follow increasing concentrations of trail pheromone |
| wiggle | turtle procedure | makes ants oscillate between left and right while walking |
| trail-pheromone-scent-at-angle | patch report procedure | reports quantity of trail pheromone at a given angle, based on "uphill-trail-pheromone" |
