## Supplementary material for "Temporary prey storage along swarm columns of army ants: an adaptive strategy for successful raiding?": Table S4

**Probability to cache/collect cached food: 25%**

| Contrast: caches vs. no caches | Estimate | Std. error | t value | p |
|---|---|---|---|---|
| brood evacuation speed 1; 50 raiders | -0.0012788 | 0.1722436 | -0.007 | 0.99408 |
| brood evacuation speed 2; 50 raiders | 0.1214595 | 0.1738161 | 0.699 | 0.48482 |
| brood evacuation speed 3; 50 raiders | 0.0196341 | 0.1742692 | 0.113 | 0.91031 |
| brood evacuation speed 4; 50 raiders | 0.0007186 | 0.182608 | 0.004 | 0.99686 |
| brood evacuation speed 5; 50 raiders | -0.1262349 | 0.1946532 | -0.649 | 0.51678 |
| brood evacuation speed 1; 100 raiders | -0.0737631 | 0.1756776 | -0.42 | 0.67465 |
| brood evacuation speed 2; 100 raiders | 0.0086986 | 0.1762047 | 0.049 | 0.96064 |
| brood evacuation speed 3; 100 raiders | 0.0500104 | 0.1795626 | 0.279 | 0.78067 |
| brood evacuation speed 4; 100 raiders | -0.0154813 | 0.1849543 | -0.084 | 0.93331 |
| brood evacuation speed 5; 100 raiders | -0.0017794 | 0.2031788 | -0.009 | 0.99301 |
| brood evacuation speed 1; 300 raiders | -0.0072098 | 0.1829052 | -0.039 | 0.96856 |
| brood evacuation speed 2; 300 raiders | 0.0125088 | 0.1795723 | 0.07 | 0.94448 |
| brood evacuation speed 3; 300 raiders | -0.2074938 | 0.1806931 | -1.148 | 0.25106 |
| brood evacuation speed 4; 300 raiders | -0.1550146 | 0.1929371 | -0.803 | 0.42188 |
| brood evacuation speed 5; 300 raiders | -0.2410106 | 0.2096095 | -1.15 | 0.25045 |
| brood evacuation speed 1; 500 raiders | 0.1673726 | 0.1719153 | 0.974 | 0.33046 |
| brood evacuation speed 2; 500 raiders | 0.1471723 | 0.1733765 | 0.849 | 0.39613 |
| brood evacuation speed 3; 500 raiders | -0.2363888 | 0.1988449 | -1.189 | 0.23475 |
| brood evacuation speed 4; 500 raiders | 0.097276 | 0.1957337 | 0.497 | 0.61929 |
| brood evacuation speed 5; 500 raiders | -0.3179169 | 0.2175375 | -1.461 | 0.14416 |
| brood evacuation speed 1; 700 raiders | -0.0302245 | 0.1436216 | -0.21 | 0.83336 |
| brood evacuation speed 2; 700 raiders | 0.7494649 | 0.1396447 | 5.367 | $p < 0.001$ |
| brood evacuation speed 3; 700 raiders | -0.2935946 | 0.1298192 | -2.262 | $p < 0.05$ |
| brood evacuation speed 4; 700 raiders | -0.423603 | 0.1423683 | -2.975 | $p < 0.01$ |
| brood evacuation speed 5; 700 raiders | -0.1470876 | 0.1696458 | -0.867 | 0.3861 |

**Probability to cache/collect cached food: 75%**

| Contrast: caches vs. no caches | Estimate | Std. error | t value | p |
|---|---|---|---|---|
| brood evacuation speed 1; 50 raiders | -0.065291 | 0.162322 | -0.402 | 0.687584 |
| brood evacuation speed 2; 50 raiders | 0.030431 | 0.176755 | 0.172 | 0.863336 |
| brood evacuation speed 3; 50 raiders | 0.047274 | 0.174682 | 0.271 | 0.786722 |
| brood evacuation speed 4; 50 raiders | 0.039459 | 0.185153 | 0.213 | 0.831274 |
| brood evacuation speed 5; 50 raiders | -0.184429 | 0.193386 | -0.954 | 0.340436 |
| brood evacuation speed 1; 100 raiders | 0.035682 | 0.174396 | 0.205 | 0.837917 |
| brood evacuation speed 2; 100 raiders | -0.190614 | 0.17944 | -1.062 | 0.288326 |
| brood evacuation speed 3; 100 raiders | 0.076414 | 0.174849 | 0.437 | 0.662169 |
| brood evacuation speed 4; 100 raiders | -0.013455 | 0.189072 | -0.071 | 0.94328 |
| brood evacuation speed 5; 100 raiders | -0.006479 | 0.204454 | -0.032 | 0.974727 |
| brood evacuation speed 1; 300 raiders | -0.147158 | 0.17908 | -0.822 | 0.411387 |
| brood evacuation speed 2; 300 raiders | -0.141678 | 0.176115 | -0.804 | 0.421289 |
| brood evacuation speed 3; 300 raiders | -0.198226 | 0.188732 | -1.05 | 0.29379 |
| brood evacuation speed 4; 300 raiders | -0.210089 | 0.186778 | -1.125 | 0.260896 |
| brood evacuation speed 5; 300 raiders | 0.021053 | 0.2128 | 0.099 | 0.921206 |
| brood evacuation speed 1; 500 raiders | -0.167304 | 0.184097 | -0.909 | 0.363647 |
| brood evacuation speed 2; 500 raiders | 0.155879 | 0.183909 | 0.848 | 0.396838 |
| brood evacuation speed 3; 500 raiders | -0.030219 | 0.192077 | -0.157 | 0.875014 |
| brood evacuation speed 4; 500 raiders | -0.128105 | 0.189832 | -0.675 | 0.499911 |
| brood evacuation speed 5; 500 raiders | 0.033902 | 0.212235 | 0.16 | 0.873116 |
| brood evacuation speed 1; 700 raiders | 0.117242 | 0.137433 | 0.853 | 0.393782 |
| brood evacuation speed 2; 700 raiders | 0.412306 | 0.137304 | 3.003 | $p < 0.01$ |
| brood evacuation speed 3; 700 raiders | -0.140582 | 0.160272 | -0.877 | 0.380584 |
| brood evacuation speed 4; 700 raiders | -0.078639 | 0.142537 | -0.552 | 0.58125 |
| brood evacuation speed 5; 700 raiders | 0.07133 | 0.18047 | 0.395 | 0.69273 |

**Probability to cache/collect cached food: 50%**

| Contrast: caches vs. no caches | Estimate | Std. error | t value | p |
|---|---|---|---|---|
| brood evacuation speed 1; 50 raiders | 7.59E-02 | 1.59E-01 | 0.476 | 0.63407 |
| brood evacuation speed 2; 50 raiders | -1.64E-16 | 1.67E-01 | 0 | 1 |
| brood evacuation speed 3; 50 raiders | 1.54E-01 | 1.73E-01 | 0.891 | 0.37306 |
| brood evacuation speed 4; 50 raiders | -8.70E-02 | 1.70E-01 | -0.512 | 0.60897 |
| brood evacuation speed 5; 50 raiders | -3.56E-02 | 1.87E-01 | -0.191 | 0.84889 |
| brood evacuation speed 1; 100 raiders | -2.82E-02 | 1.70E-01 | -0.166 | 0.86834 |
| brood evacuation speed 2; 100 raiders | -2.28E-01 | 1.72E-01 | -1.328 | 0.18456 |
| brood evacuation speed 3; 100 raiders | -6.43E-02 | 1.77E-01 | -0.364 | 0.71578 |
| brood evacuation speed 4; 100 raiders | -3.06E-02 | 1.82E-01 | -0.169 | 0.86618 |
| brood evacuation speed 5; 100 raiders | -1.21E-01 | 1.99E-01 | -0.604 | 0.54576 |
| brood evacuation speed 1; 300 raiders | -7.17E-02 | 1.81E-01 | -0.397 | 0.69123 |
| brood evacuation speed 2; 300 raiders | -7.43E-02 | 1.77E-01 | -0.419 | 0.67521 |
| brood evacuation speed 3; 300 raiders | -2.25E-02 | 1.74E-01 | -0.129 | 0.89731 |
| brood evacuation speed 4; 300 raiders | -7.81E-02 | 1.84E-01 | -0.425 | 0.67064 |
| brood evacuation speed 5; 300 raiders | -5.45E-02 | 2.09E-01 | -0.261 | 0.79399 |
| brood evacuation speed 1; 500 raiders | -7.17E-02 | 1.71E-01 | -0.42 | 0.67482 |
| brood evacuation speed 2; 500 raiders | 4.54E-02 | 1.89E-01 | 0.241 | 0.80982 |
| brood evacuation speed 3; 500 raiders | -4.16E-02 | 1.69E-01 | -0.246 | 0.80546 |
| brood evacuation speed 4; 500 raiders | 6.67E-02 | 1.80E-01 | 0.371 | 0.71085 |
| brood evacuation speed 5; 500 raiders | -1.41E-01 | 1.99E-01 | -0.708 | 0.479 |
| brood evacuation speed 1; 700 raiders | 1.74E-01 | 1.26E-01 | 1.379 | 0.16823 |
| brood evacuation speed 2; 700 raiders | -1.15E-01 | 1.38E-01 | -0.831 | 0.40586 |
| brood evacuation speed 3; 700 raiders | 3.00E-01 | 1.35E-01 | 2.232 | $p < 0.05$ |
| brood evacuation speed 4; 700 raiders | 3.66E-01 | 1.40E-01 | 2.62 | $p < 0.01$ |
| brood evacuation speed 5; 700 raiders | 7.79E-02 | 1.78E-01 | 0.437 | 0.66198 |

**Probability to cache/collect cached food: 100%**

| Contrast: caches vs. no caches | Estimate | Std. error | t value | p |
|---|---|---|---|---|
| brood evacuation speed 1; 50 raiders | -0.146731 | 0.189528 | -0.774 | 0.43897 |
| brood evacuation speed 2; 50 raiders | -0.123984 | 0.189414 | -0.655 | 0.51287 |
| brood evacuation speed 3; 50 raiders | -0.178016 | 0.195929 | -0.909 | 0.36376 |
| brood evacuation speed 4; 50 raiders | -0.067695 | 0.20371 | -0.332 | 0.73971 |
| brood evacuation speed 5; 50 raiders | 0.093511 | 0.212806 | 0.439 | 0.66044 |
| brood evacuation speed 1; 100 raiders | -0.13783 | 0.184882 | -0.745 | 0.45612 |
| brood evacuation speed 2; 100 raiders | -0.016218 | 0.189118 | -0.086 | 0.93167 |
| brood evacuation speed 3; 100 raiders | -0.009472 | 0.193133 | -0.049 | 0.96089 |
| brood evacuation speed 4; 100 raiders | -0.005944 | 0.202381 | -0.029 | 0.97658 |
| brood evacuation speed 5; 100 raiders | -0.198765 | 0.229181 | -0.867 | 0.38596 |
| brood evacuation speed 1; 300 raiders | 0.015279 | 0.200276 | 0.076 | 0.9392 |
| brood evacuation speed 2; 300 raiders | 0.128072 | 0.210354 | 0.609 | 0.54274 |
| brood evacuation speed 3; 300 raiders | 0.165181 | 0.218276 | 0.757 | 0.44935 |
| brood evacuation speed 4; 300 raiders | -0.88584 | 0.182647 | -4.85 | $p < 0.001$ |
| brood evacuation speed 5; 300 raiders | 0.016461 | 0.238161 | 0.069 | 0.94491 |
| brood evacuation speed 1; 500 raiders | 0.057021 | 0.210472 | 0.271 | 0.7865 |
| brood evacuation speed 2; 500 raiders | -0.341986 | 0.205997 | -1.66 | 0.09715 |
| brood evacuation speed 3; 500 raiders | -0.077725 | 0.213612 | -0.364 | 0.71603 |
| brood evacuation speed 4; 500 raiders | -0.033351 | 0.217145 | -0.154 | 0.87796 |
| brood evacuation speed 5; 500 raiders | -0.075473 | 0.225364 | -0.335 | 0.73776 |
| brood evacuation speed 1; 700 raiders | -0.262293 | 0.149775 | -1.751 | 0.08016 |
| brood evacuation speed 2; 700 raiders | 0.138966 | 0.166572 | 0.834 | 0.4043 |
| brood evacuation speed 3; 700 raiders | -0.197535 | 0.150799 | -1.31 | 0.19047 |
| brood evacuation speed 4; 700 raiders | 0.285447 | 0.164147 | 1.739 | 0.0823 |
| brood evacuation speed 5; 700 raiders | -0.137539 | 0.164815 | -0.835 | 0.40416 |
