## Supplementary material for "Temporary prey storage along swarm columns of army ants: an adaptive strategy for successful raiding?": Table S5

**Probability to cache/collect cached food: 25%**

| Contrast: continuous vs. single caching | Estimate | Std. error | t value | p |
|---|---|---|---|---|
| brood evacuation speed 1; 50 raiders | -0.021715 | 0.039931 | -0.544 | 0.586669 |
| brood evacuation speed 2; 50 raiders | -0.014072 | 0.041316 | -0.341 | 0.733474 |
| brood evacuation speed 3; 50 raiders | 0.040375 | 0.043701 | 0.924 | 0.355731 |
| brood evacuation speed 4; 50 raiders | -0.047356 | 0.047737 | -0.992 | 0.321394 |
| brood evacuation speed 5; 50 raiders | -0.024553 | 0.072595 | -0.338 | 0.735259 |
| brood evacuation speed 1; 100 raiders | 0.010074 | 0.036585 | 0.275 | 0.783082 |
| brood evacuation speed 2; 100 raiders | -0.01754 | 0.03851 | -0.455 | 0.648859 |
| brood evacuation speed 3; 100 raiders | -0.014193 | 0.04062 | -0.349 | 0.726845 |
| brood evacuation speed 4; 100 raiders | -0.029446 | 0.044767 | -0.658 | 0.51082 |
| brood evacuation speed 5; 100 raiders | -0.011472 | 0.07304 | -0.157 | 0.875216 |
| brood evacuation speed 1; 300 raiders | -0.031615 | 0.034297 | -0.922 | 0.356819 |
| brood evacuation speed 2; 300 raiders | 0.038806 | 0.03508 | 1.106 | 0.268866 |
| brood evacuation speed 3; 300 raiders | -0.017608 | 0.038013 | -0.463 | 0.643298 |
| brood evacuation speed 4; 300 raiders | -0.04142 | 0.042171 | -0.982 | 0.326202 |
| brood evacuation speed 5; 300 raiders | 0.02725 | 0.071198 | 0.383 | 0.701986 |
| brood evacuation speed 1; 500 raiders | 0.024828 | 0.034266 | 0.725 | 0.46887 |
| brood evacuation speed 2; 500 raiders | 0.012538 | 0.035995 | 0.348 | 0.727658 |
| brood evacuation speed 3; 500 raiders | -0.071875 | 0.038416 | -1.871 | 0.061593 |
| brood evacuation speed 4; 500 raiders | 0.103042 | 0.043818 | 2.352 | < 0.05 |
| brood evacuation speed 5; 500 raiders | -0.076076 | 0.070291 | -1.082 | 0.279342 |
| brood evacuation speed 1; 700 raiders | 0.032302 | 0.034439 | 0.938 | 0.348459 |
| brood evacuation speed 2; 700 raiders | 0.007757 | 0.035681 | 0.217 | 0.827928 |
| brood evacuation speed 3; 700 raiders | 0.005423 | 0.037087 | 0.146 | 0.883774 |
| brood evacuation speed 4; 700 raiders | -0.019943 | 0.042367 | -0.471 | 0.637932 |
| brood evacuation speed 5; 700 raiders | -0.075807 | 0.070167 | -1.08 | 0.280189 |

**Probability to cache/collect cached food: 50%**

| Contrast: continuous vs. single caching | Estimate | Std. error | t value | p |
|---|---|---|---|---|
| brood evacuation speed 1; 50 raiders | 0.05984 | 0.038576 | 1.551 | 0.121111 |
| brood evacuation speed 2; 50 raiders | 0.054412 | 0.039971 | 1.361 | 0.173678 |
| brood evacuation speed 3; 50 raiders | 0.01216 | 0.042019 | 0.289 | 0.77233 |
| brood evacuation speed 4; 50 raiders | -0.00989 | 0.045812 | -0.216 | 0.829119 |
| brood evacuation speed 5; 50 raiders | 0.018349 | 0.07115 | 0.258 | 0.796532 |
| brood evacuation speed 1; 100 raiders | 0.028287 | 0.035479 | 0.797 | 0.425453 |
| brood evacuation speed 2; 100 raiders | -0.004876 | 0.036675 | -0.133 | 0.894261 |
| brood evacuation speed 3; 100 raiders | 0.030516 | 0.039126 | 0.78 | 0.435586 |
| brood evacuation speed 4; 100 raiders | 0.004813 | 0.043551 | 0.111 | 0.912026 |
| brood evacuation speed 5; 100 raiders | -0.009001 | 0.070472 | -0.128 | 0.898388 |
| brood evacuation speed 1; 300 raiders | -0.039016 | 0.033315 | -1.171 | 0.241787 |
| brood evacuation speed 2; 300 raiders | -0.022959 | 0.034902 | -0.658 | 0.51078 |
| brood evacuation speed 3; 300 raiders | 0.037218 | 0.036285 | 1.026 | 0.305238 |
| brood evacuation speed 4; 300 raiders | -0.044395 | 0.041835 | -1.061 | 0.288816 |
| brood evacuation speed 5; 300 raiders | -0.017858 | 0.07019 | -0.254 | 0.799216 |
| brood evacuation speed 1; 500 raiders | 0.062494 | 0.034141 | 1.83 | 0.067432 |
| brood evacuation speed 2; 500 raiders | 0.078767 | 0.035459 | 2.221 | < 0.05 |
| brood evacuation speed 3; 500 raiders | -0.028761 | 0.036674 | -0.784 | 0.433054 |
| brood evacuation speed 4; 500 raiders | -0.108061 | 0.041552 | -2.601 | < 0.01 |
| brood evacuation speed 5; 500 raiders | 0.025456 | 0.070828 | 0.359 | 0.719356 |
| brood evacuation speed 1; 700 raiders | 0.049876 | 0.033729 | 1.479 | 0.139467 |
| brood evacuation speed 2; 700 raiders | 0.05148 | 0.035459 | 1.452 | 0.146814 |
| brood evacuation speed 3; 700 raiders | 0.048328 | 0.037081 | 1.303 | 0.192717 |
| brood evacuation speed 4; 700 raiders | 0.036757 | 0.042942 | 0.856 | 0.392197 |
| brood evacuation speed 5; 700 raiders | -0.032203 | 0.070259 | -0.458 | 0.646785 |

**Probability to cache/collect cached food: 75%**

| Contrast: continuous vs. single caching | Estimate | Std. error | t value | p |
|---|---|---|---|---|
| brood evacuation speed 1; 50 raiders | 0.094719 | 0.038033 | 2.49 | < 0.05 |
| brood evacuation speed 2; 50 raiders | 0.088775 | 0.039094 | 2.271 | < 0.05 |
| brood evacuation speed 3; 50 raiders | 0.074157 | 0.041021 | 1.808 | 0.070891 |
| brood evacuation speed 4; 50 raiders | 0.06197 | 0.044285 | 1.399 | 0.161968 |
| brood evacuation speed 5; 50 raiders | 0.010714 | 0.067637 | 0.158 | 0.87416 |
| brood evacuation speed 1; 100 raiders | -0.008387 | 0.033627 | -0.249 | 0.803093 |
| brood evacuation speed 2; 100 raiders | 0.069912 | 0.034919 | 2.002 | < 0.05 |
| brood evacuation speed 3; 100 raiders | 0.026964 | 0.036805 | 0.733 | 0.463928 |
| brood evacuation speed 4; 100 raiders | -0.006579 | 0.041054 | -0.16 | 0.872708 |
| brood evacuation speed 5; 100 raiders | 0.007313 | 0.068435 | 0.107 | 0.914922 |
| brood evacuation speed 1; 300 raiders | -0.009169 | 0.031957 | -0.287 | 0.774229 |
| brood evacuation speed 2; 300 raiders | 0.012056 | 0.032634 | 0.369 | 0.711879 |
| brood evacuation speed 3; 300 raiders | 0.049937 | 0.035422 | 1.41 | 0.158865 |
| brood evacuation speed 4; 300 raiders | 0.02895 | 0.040599 | 0.713 | 0.475942 |
| brood evacuation speed 5; 300 raiders | -0.010526 | 0.067041 | -0.157 | 0.875259 |
| brood evacuation speed 1; 500 raiders | -0.016306 | 0.03192 | -0.511 | 0.609559 |
| brood evacuation speed 2; 500 raiders | 0.003403 | 0.03301 | 0.103 | 0.917913 |
| brood evacuation speed 3; 500 raiders | 0.048346 | 0.035025 | 1.38 | 0.167738 |
| brood evacuation speed 4; 500 raiders | -0.066185 | 0.039454 | -1.678 | 0.093704 |
| brood evacuation speed 5; 500 raiders | -0.007194 | 0.067879 | -0.106 | 0.915611 |
| brood evacuation speed 1; 700 raiders | 0.011547 | 0.032503 | 0.355 | 0.722468 |
| brood evacuation speed 2; 700 raiders | 0.064364 | 0.033968 | 1.895 | 0.058355 |
| brood evacuation speed 3; 700 raiders | 0.005431 | 0.035564 | 0.153 | 0.878656 |
| brood evacuation speed 4; 700 raiders | -0.008279 | 0.038923 | -0.213 | 0.831591 |
| brood evacuation speed 5; 700 raiders | 0.108308 | 0.066962 | 1.617 | 0.106043 |

**Probability to cache/collect cached food: 100%**

| Contrast: continuous vs. single caching | Estimate | Std. error | t value | p |
|---|---|---|---|---|
| brood evacuation speed 1; 50 raiders | -2.34E-01 | 4.62E-02 | -5.059 | < 0.001 |
| brood evacuation speed 2; 50 raiders | -6.10E-02 | 5.08E-02 | -1.201 | 0.230083 |
| brood evacuation speed 3; 50 raiders | -2.10E-01 | 4.96E-02 | -4.24 | < 0.001 |
| brood evacuation speed 4; 50 raiders | -1.96E-01 | 5.23E-02 | -3.75 | < 0.001 |
| brood evacuation speed 5; 50 raiders | -3.48E-02 | 7.72E-02 | -0.45 | 0.65277 |
| brood evacuation speed 1; 100 raiders | -9.88E-02 | 3.89E-02 | -2.54 | < 0.05 |
| brood evacuation speed 2; 100 raiders | -9.27E-02 | 3.98E-02 | -2.329 | < 0.05 |
| brood evacuation speed 3; 100 raiders | -1.26E-01 | 4.22E-02 | -2.973 | < 0.01 |
| brood evacuation speed 4; 100 raiders | -5.65E-02 | 4.70E-02 | -1.202 | 0.229446 |
| brood evacuation speed 5; 100 raiders | 2.11E-02 | 7.70E-02 | 0.274 | 0.783851 |
| brood evacuation speed 1; 300 raiders | 8.18E-04 | 3.55E-02 | 0.023 | 0.981641 |
| brood evacuation speed 2; 300 raiders | -1.72E-02 | 3.65E-02 | -0.472 | 0.636751 |
| brood evacuation speed 3; 300 raiders | -1.69E-02 | 3.92E-02 | -0.432 | 0.666146 |
| brood evacuation speed 4; 300 raiders | -4.70E-02 | 4.21E-02 | -1.117 | 0.264242 |
| brood evacuation speed 5; 300 raiders | 2.62E-02 | 7.61E-02 | 0.345 | 0.730345 |
| brood evacuation speed 1; 500 raiders | -2.24E-02 | 3.55E-02 | -0.632 | 0.527772 |
| brood evacuation speed 2; 500 raiders | -2.27E-02 | 3.74E-02 | -0.606 | 0.544594 |
| brood evacuation speed 3; 500 raiders | 3.33E-02 | 3.98E-02 | 0.837 | 0.402609 |
| brood evacuation speed 4; 500 raiders | -1.13E-01 | 4.47E-02 | -2.527 | < 0.05 |
| brood evacuation speed 5; 500 raiders | -1.05E-02 | 7.35E-02 | -0.143 | 0.886405 |
| brood evacuation speed 1; 700 raiders | 1.01E-01 | 3.62E-02 | 2.799 | < 0.01 |
| brood evacuation speed 2; 700 raiders | 1.55E-02 | 3.70E-02 | 0.42 | 0.674885 |
| brood evacuation speed 3; 700 raiders | 1.05E-01 | 3.99E-02 | 2.621 | < 0.01 |
| brood evacuation speed 4; 700 raiders | 1.00E-01 | 4.57E-02 | 2.188 | < 0.05 |
| brood evacuation speed 5; 700 raiders | -1.78E-03 | 7.40E-02 | -0.024 | 0.980879 |
