## Supplementary material for "Temporary prey storage along swarm columns of army ants: an adaptive strategy for successful raiding?": Table S6

**Probability to cache/collect cached food: 25%**

| Contrast: continuous vs. single caching | Estimate | Std. error | t value | p |
|---|---|---|---|---|
| brood evacuation speed 1; 50 raiders | -0.81554 | 0.154079 | -5.293 | < 0.001 |
| brood evacuation speed 2; 50 raiders | -0.75107 | 0.152179 | -4.935 | < 0.001 |
| brood evacuation speed 3; 50 raiders | -0.813591 | 0.155121 | -5.245 | < 0.001 |
| brood evacuation speed 4; 50 raiders | -0.828025 | 0.162958 | -5.081 | < 0.001 |
| brood evacuation speed 5; 50 raiders | -0.669431 | 0.183751 | -3.643 | < 0.001 |
| brood evacuation speed 1; 100 raiders | -0.495957 | 0.168956 | -2.935 | < 0.01 |
| brood evacuation speed 2; 100 raiders | -0.475982 | 0.166693 | -2.855 | < 0.01 |
| brood evacuation speed 3; 100 raiders | -0.42868 | 0.169652 | -2.527 | < 0.05 |
| brood evacuation speed 4; 100 raiders | -0.369152 | 0.179792 | -2.053 | < 0.05 |
| brood evacuation speed 5; 100 raiders | -0.343318 | 0.197885 | -1.735 | 0.083009 |
| brood evacuation speed 1; 300 raiders | -0.00721 | 0.192602 | -0.037 | 0.970145 |
| brood evacuation speed 2; 300 raiders | 0.001382 | 0.188565 | 0.007 | 0.994153 |
| brood evacuation speed 3; 300 raiders | 0.022773 | 0.201013 | 0.113 | 0.909819 |
| brood evacuation speed 4; 300 raiders | -0.153533 | 0.203235 | -0.755 | 0.450129 |
| brood evacuation speed 5; 300 raiders | -0.03029 | 0.231833 | -0.131 | 0.896072 |
| brood evacuation speed 1; 500 raiders | 0.219065 | 0.183612 | 1.193 | 0.233071 |
| brood evacuation speed 2; 500 raiders | 0.108968 | 0.180722 | 0.603 | 0.546648 |
| brood evacuation speed 3; 500 raiders | -0.326216 | 0.20538 | -1.588 | 0.11247 |
| brood evacuation speed 4; 500 raiders | -0.275034 | 0.188567 | -1.459 | 0.144952 |
| brood evacuation speed 5; 500 raiders | -0.270415 | 0.231405 | -1.169 | 0.242806 |
| brood evacuation speed 1; 700 raiders | -0.145084 | 0.147139 | -0.986 | 0.324315 |
| brood evacuation speed 2; 700 raiders | 0.444384 | 0.133275 | 3.334 | < 0.001 |
| brood evacuation speed 3; 700 raiders | 0.229091 | 0.155458 | 1.474 | 0.140837 |
| brood evacuation speed 4; 700 raiders | -0.303657 | 0.153648 | -1.976 | < 0.05 |
| brood evacuation speed 5; 700 raiders | -0.195878 | 0.176657 | -1.109 | 0.267738 |

**Probability to cache/collect cached food: 75%**

| Contrast: continuous vs. single caching | Estimate | Std. error | t value | p |
|---|---|---|---|---|
| brood evacuation speed 1; 50 raiders | -0.79962 | 0.17962 | -4.452 | < 0.001 |
| brood evacuation speed 2; 50 raiders | -0.90506 | 0.18801 | -4.814 | < 0.001 |
| brood evacuation speed 3; 50 raiders | -0.7878 | 0.1883 | -4.184 | < 0.001 |
| brood evacuation speed 4; 50 raiders | -0.77044 | 0.20053 | -3.842 | < 0.001 |
| brood evacuation speed 5; 50 raiders | -0.74812 | 0.22185 | -3.372 | < 0.001 |
| brood evacuation speed 1; 100 raiders | -0.42986 | 0.20083 | -2.14 | < 0.05 |
| brood evacuation speed 2; 100 raiders | -0.6144 | 0.21082 | -2.914 | < 0.01 |
| brood evacuation speed 3; 100 raiders | -0.43117 | 0.19921 | -2.164 | < 0.05 |
| brood evacuation speed 4; 100 raiders | -0.55813 | 0.21515 | -2.594 | < 0.01 |
| brood evacuation speed 5; 100 raiders | -0.44936 | 0.23705 | -1.896 | 0.058248 |
| brood evacuation speed 1; 300 raiders | -0.13057 | 0.22994 | -0.568 | 0.570245 |
| brood evacuation speed 2; 300 raiders | -0.14232 | 0.22523 | -0.632 | 0.527598 |
| brood evacuation speed 3; 300 raiders | -0.22002 | 0.24023 | -0.916 | 0.359918 |
| brood evacuation speed 4; 300 raiders | -0.15098 | 0.24214 | -0.624 | 0.533069 |
| brood evacuation speed 5; 300 raiders | -0.07545 | 0.26579 | -0.284 | 0.776548 |
| brood evacuation speed 1; 500 raiders | -0.3324 | 0.22712 | -1.464 | 0.143575 |
| brood evacuation speed 2; 500 raiders | 0.03086 | 0.22767 | 0.136 | 0.892207 |
| brood evacuation speed 3; 500 raiders | -0.25994 | 0.23294 | -1.116 | 0.264682 |
| brood evacuation speed 4; 500 raiders | -0.07599 | 0.24583 | -0.309 | 0.757302 |
| brood evacuation speed 5; 500 raiders | 0.05647 | 0.27304 | 0.207 | 0.836194 |
| brood evacuation speed 1; 700 raiders | 0.08758 | 0.17442 | 0.502 | 0.615665 |
| brood evacuation speed 2; 700 raiders | 0.46755 | 0.1786 | 2.618 | < 0.01 |
| brood evacuation speed 3; 700 raiders | -0.43721 | 0.19238 | -2.273 | < 0.05 |
| brood evacuation speed 4; 700 raiders | -0.27533 | 0.17432 | -1.579 | 0.114499 |
| brood evacuation speed 5; 700 raiders | -0.73135 | 0.19509 | -3.749 | < 0.001 |

**Probability to cache/collect cached food: 50%**

| Contrast: continuous vs. single caching | Estimate | Std. error | t value | p |
|---|---|---|---|---|
| brood evacuation speed 1; 50 raiders | -0.899388 | 0.153338 | -5.865 | < 0.001 |
| brood evacuation speed 2; 50 raiders | -0.955017 | 0.162127 | -5.891 | < 0.001 |
| brood evacuation speed 3; 50 raiders | -0.883339 | 0.163478 | -5.403 | < 0.001 |
| brood evacuation speed 4; 50 raiders | -0.859206 | 0.171351 | -5.014 | < 0.001 |
| brood evacuation speed 5; 50 raiders | -0.644625 | 0.192545 | -3.348 | < 0.001 |
| brood evacuation speed 1; 100 raiders | -0.612252 | 0.176146 | -3.476 | < 0.001 |
| brood evacuation speed 2; 100 raiders | -0.706784 | 0.183353 | -3.855 | < 0.001 |
| brood evacuation speed 3; 100 raiders | -0.614782 | 0.184222 | -3.337 | < 0.001 |
| brood evacuation speed 4; 100 raiders | -0.603729 | 0.188417 | -3.204 | < 0.01 |
| brood evacuation speed 5; 100 raiders | -0.407093 | 0.219216 | -1.857 | 0.06355 |
| brood evacuation speed 1; 300 raiders | -0.151464 | 0.20723 | -0.731 | 0.464984 |
| brood evacuation speed 2; 300 raiders | -0.092418 | 0.206597 | -0.447 | 0.654715 |
| brood evacuation speed 3; 300 raiders | -0.031438 | 0.203095 | -0.155 | 0.877009 |
| brood evacuation speed 4; 300 raiders | -0.06601 | 0.215459 | -0.306 | 0.759377 |
| brood evacuation speed 5; 300 raiders | -0.188526 | 0.236591 | -0.797 | 0.425698 |
| brood evacuation speed 1; 500 raiders | -0.118873 | 0.197678 | -0.601 | 0.547722 |
| brood evacuation speed 2; 500 raiders | 0.007296 | 0.218764 | 0.033 | 0.973399 |
| brood evacuation speed 3; 500 raiders | 0.114002 | 0.205309 | 0.555 | 0.578813 |
| brood evacuation speed 4; 500 raiders | -0.028573 | 0.205363 | -0.139 | 0.889366 |
| brood evacuation speed 5; 500 raiders | 0.038043 | 0.24313 | 0.156 | 0.875688 |
| brood evacuation speed 1; 700 raiders | 0.27481 | 0.151999 | 1.808 | 0.070861 |
| brood evacuation speed 2; 700 raiders | 0.046498 | 0.168248 | 0.276 | 0.782316 |
| brood evacuation speed 3; 700 raiders | -0.238887 | 0.137317 | -1.74 | 0.082173 |
| brood evacuation speed 4; 700 raiders | 0.024767 | 0.148841 | 0.166 | 0.867873 |
| brood evacuation speed 5; 700 raiders | -0.147567 | 0.197121 | -0.749 | 0.454239 |

**Probability to cache/collect cached food: 100%**

| Contrast: continuous vs. single caching | Estimate | Std. error | t value | p |
|---|---|---|---|---|
| brood evacuation speed 1; 50 raiders | -0.1382406 | 0.2129947 | -0.649 | 0.51644 |
| brood evacuation speed 2; 50 raiders | 0.011818 | 0.2195737 | 0.054 | 0.95709 |
| brood evacuation speed 3; 50 raiders | -0.0320803 | 0.2274837 | -0.141 | 0.88788 |
| brood evacuation speed 4; 50 raiders | -0.0139106 | 0.2315107 | -0.06 | 0.9521 |
| brood evacuation speed 5; 50 raiders | 0.0237729 | 0.2344384 | 0.101 | 0.91925 |
| brood evacuation speed 1; 100 raiders | -0.0019861 | 0.2142731 | -0.009 | 0.99261 |
| brood evacuation speed 2; 100 raiders | 0.0867141 | 0.2177424 | 0.398 | 0.69052 |
| brood evacuation speed 3; 100 raiders | 0.1060406 | 0.2231206 | 0.475 | 0.63469 |
| brood evacuation speed 4; 100 raiders | -0.0096404 | 0.2267813 | -0.043 | 0.9661 |
| brood evacuation speed 5; 100 raiders | 0.0126451 | 0.2703357 | 0.047 | 0.9627 |
| brood evacuation speed 1; 300 raiders | 0.1268493 | 0.2312128 | 0.549 | 0.58336 |
| brood evacuation speed 2; 300 raiders | 0.183374 | 0.2394748 | 0.766 | 0.44399 |
| brood evacuation speed 3; 300 raiders | 0.2199765 | 0.2485219 | 0.885 | 0.37626 |
| brood evacuation speed 4; 300 raiders | -1.3237533 | 0.1939622 | -6.825 | < 0.001 |
| brood evacuation speed 5; 300 raiders | 0.0907289 | 0.2722631 | 0.333 | 0.73901 |
| brood evacuation speed 1; 500 raiders | -0.24339 | 0.2197513 | -1.108 | 0.26827 |
| brood evacuation speed 2; 500 raiders | -0.1693363 | 0.2399179 | -0.706 | 0.48044 |
| brood evacuation speed 3; 500 raiders | -0.085649 | 0.239132 | -0.358 | 0.72028 |
| brood evacuation speed 4; 500 raiders | -0.0155309 | 0.244624 | -0.063 | 0.94939 |
| brood evacuation speed 5; 500 raiders | 0.1492234 | 0.2676502 | 0.558 | 0.57727 |
| brood evacuation speed 1; 700 raiders | -0.0177803 | 0.1778164 | -0.1 | 0.92037 |
| brood evacuation speed 2; 700 raiders | -0.0009363 | 0.1801885 | -0.005 | 0.99585 |
| brood evacuation speed 3; 700 raiders | -0.145635 | 0.1711547 | -0.851 | 0.395 |
| brood evacuation speed 4; 700 raiders | -0.0499081 | 0.1684703 | -0.296 | 0.7671 |
| brood evacuation speed 5; 700 raiders | -0.0508192 | 0.1887119 | -0.269 | 0.78775 |
