## Supplementary material for "Temporary prey storage along swarm columns of army ants: an adaptive strategy for successful raiding?": Table S7

| Model | Response variable | Factors | Model distribution |
|---|---|---|---|
| GLMs (4) | retrieved prey item number | using/not using caches nested in raid size nested in brood evacuation speed, for all probabilities of caching/collecting cached prey items | Quasipoisson |
| GLMs (4) | time spent retrieving all collected prey | using/not using caches nested in raid size nested in brood evacuation speed, for all probabilities of caching/collecting cached prey items | Quasipoisson |
| GLM | retrieved prey item number | brood evacuation speed, raider numbers, probability to cache/collect cached prey items | Quasipoisson |
| GLM | time spent retrieving all collected prey | brood evacuation speed, raider numbers, probability to cache/collect cached prey items | Quasipoisson |
